# Cell type and regulatory analysis in amphioxus illuminates evolutionary origin of the vertebrate head

**DOI:** 10.1101/2024.01.18.576194

**Authors:** Anna Markos, Jan Kubovciak, Simona Mikula Mrstakova, Anna Zitova, Jan Paces, Simona Machacova, Zbynek Kozmik, Zbynek Kozmik, Iryna Kozmikova

## Abstract

To shed light on the enigmatic origin of the vertebrate head, our study employs an integrated approach that combines single-cell transcriptomics, perturbations in signalling pathways, and cis-regulatory analysis in amphioxus, a close relative of chordate common ancestor. Through cell type characterization, we identified the presence of a prechordal plate, pre-migratory and migratory neural crest-like cell populations in the developing amphioxus embryo. Functional analysis established conserved roles of the Nodal and Hedgehog signalling pathways in prechordal plate, and of Wnt signalling pathway in neural crest development. Furthermore, the trans-species transgenic experiments provided evidence of cis-regulatory level homology within the chordate lineage. Our findings provide evidence that the key features of vertebrate head development can be traced back to the common ancestor of all chordates.

**One Sentence Summary:** Cell populations forming the vertebrate head are present in the close relative of chordate common ancestor.

## Main Text

Deciphering the enigmatic evolution and origin of the vertebrate head stands as a pivotal challenge in the realm of chordate evolutionary developmental biology. Central to this enigma are the neural crest (NC), placodes, and prechordal plate (PrCP), innovations fundamental to vertebrate head evolution, yet intriguingly absent in amphioxus, the closest living relative to chordate common ancestor (*1–5*).

The PrCP is a transient embryonic structure that forms as a band of thickened mesendoderm, rostrally to the notochord (*6–9*). It significantly influences the induction and patterning of the forebrain (*10, 11*) and contributes to the cranial mesoderm and endoderm (*12, 13*). Based on single-cell RNA sequencing (scRNA-seq) analysis in zebrafish, a transcriptional divergence between the PrCP and the notochord emerges during gastrulation, leading to the formation of two distinct cell populations with specific transcriptomic profiles (*14*).

Several studies demonstrated the existence of a gastrula organizer in amphioxus, with a compelling degree of homology to the dorsal organizer of vertebrates (*15–18*). Despite this, a prevailing view suggests the absence of a PrCP in amphioxus (*4, 19*). This assertion stems primarily from two distinctive features observed in amphioxus - the extension of the notochord beyond the brain vesicle in adults, and the inconsistent gene expression patterns that defy the conventional model of PrCP formation in vertebrates (*4, 19, 20*).

A recent study revealed that the cerebral vesicle of amphioxus at the early mid-neurula stage displays vertebrate-like partitioning, including an area termed the hypothalamo-prethalamic primordium (HyPth), which exhibits molecular partitions similar to those of the vertebrate floor, basal, and alar/roof plates (*21*). In vertebrates, these partitions are influenced by signals from the PrCP(*10*). The work on HyPth (*21*) has revived the debate on the existence of a PrCP in amphioxus. However, despite generating new hypotheses, the question remained unanswered due to lack of conclusive experimental data (*20, 22*).

Here, to address the question of head evolution in the chordate lineage, we utilized a single-cell transcriptomic approach to analyze embryonic cells across four developmental stages in amphioxus. Initially, we identified a PrCP population, as well as pre-migratory and migratory neural crest-like populations in the amphioxus head region. These populations are characterized by a unique combination of differentially expressed genes (DEGs), the orthologs of which are found in the vertebrate PrCP (*14*), as well as in neural crest (NC) or its derivatives (*23–30*). Nodal and Hh signalling are fundamental to PrCP and head development in vertebrates (*7, 9, 31*), and Wnt signaling is required for NC specification (*32*). Therefore, we examined the roles of the Nodal and Hh (Hedgehog) signalling pathways in the development of amphioxus PrCP, along with Wnt signaling in neural crest-like populations. Trans-species transgenic experiments further provided evidence of the conservation of PrCP and NCC developmental programs within the chordate lineage at the cis-regulatory level.

## Results

### PrCP is present in amphioxus embryo

To determine the existence of a PrCP in amphioxus, we clustered scRNA-seq datasets from mid-gastrula to neurula stages (G4, N0, N2 and N5) and annotated the resulting clusters based on the spatial expression patterns of marker genes (Fig.1; fig.S1, S1a-d, S2, S*2*a-e, S3, S3a-e, S4, S4a-e).

**Fig.1:**
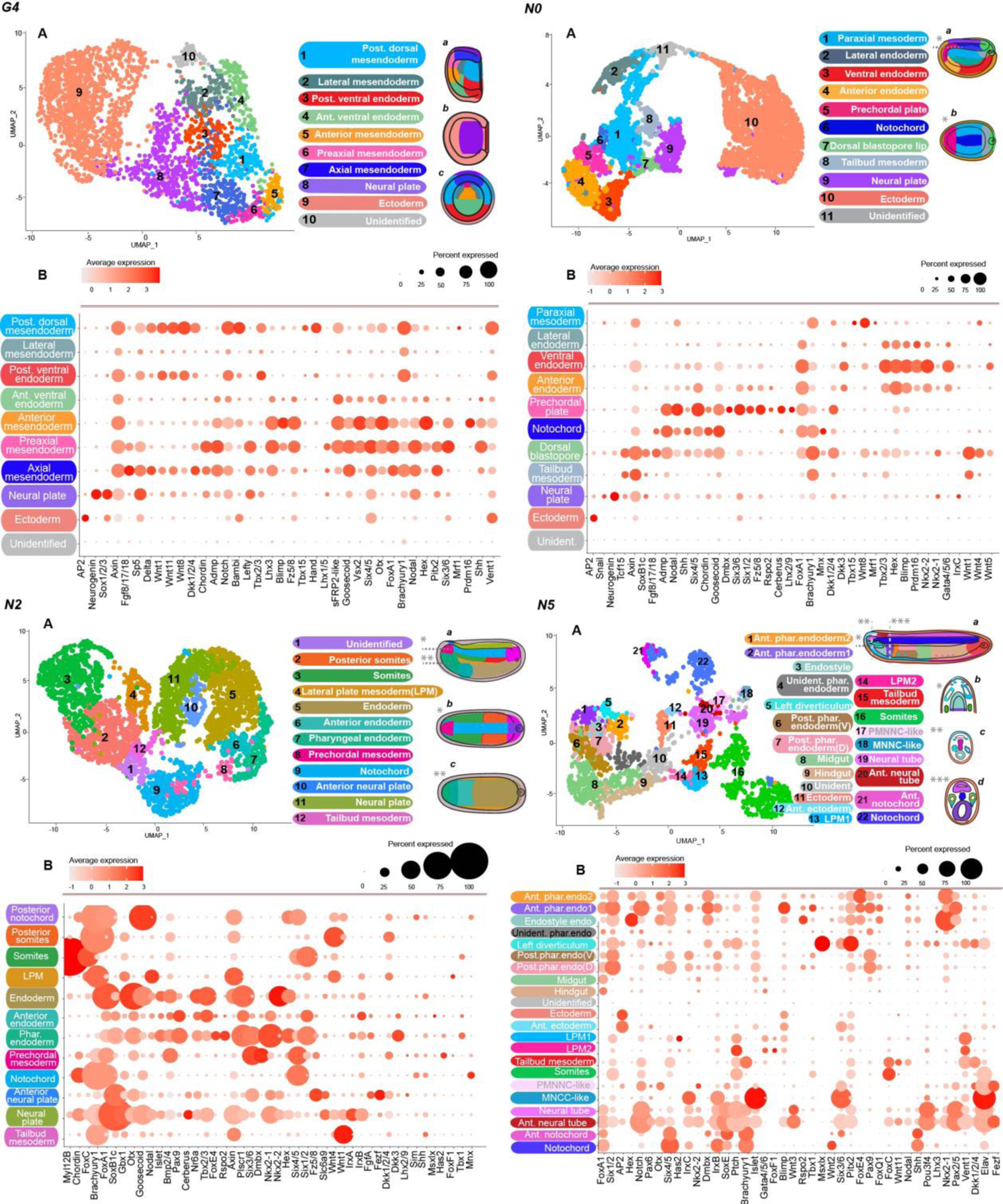
Identification of cell populations at the G4, N0, N2, and N5 stages of amphioxus with scRNA-seq analysis. (**A**) The UMAP plots of scRNA-seq clustering. (**B**) The DotPlots depict the expression of marker genes for the identified cell populations. (***a-d***) Schematic drawings illustrate the spatial position of individual clusters in the context of the amphioxus embryo. (***a***) Lateral view. (***G4b***) Dorsal view. (***G4c***) Blastopore view. The sections (***N0b, N2bc, N5b-d***) through ***G4a***, ***N2a***, ***N5a*** are indicated with dashed lines. Post.-posterior. Ant.-anterior. Phar.-pharyngeal. Endo – endoderm. Unident.-unidentified. (V) – ventral, (D) – dorsal. PMNCC-like – pre-migratory neural crest cells-like. MNCC-like – migratory neural crest cells-like.

Our analysis primarily targeted clusters in the axial and anterior mesendoderm territory. At G4, we identified axial, preaxial and anterior mesendoderm cell populations (Fig. 1*G4*A-B). The preaxial mesendoderm is characterized by the abundant expression of Chordin, Shh, Six3/6, Six4/5, Nodal, Goosecoid, Pitx2 and Lefty genes. While axial and anterior mesendoderm share small number of common genes between each other, preaxial mesendoderm exhibits a higher degree of overlap in expressed genes with both axial and anterior mesendoderm populations.

At N0, we identified a cluster expressing Lhx2/9 and Dmbx (Fig.1*N0*A-B: **fig.S2D1-E1**), along with predominant expression of Six3/6, Nodal, Fz5/8, Cerberus, and Dkk1/2/4, all recognized as PrCP markers in vertebrates (*14, 33*). These cells, representing the anterior dorsal mesendoderm domain (Fig.1*N0*Aa-b), contrast the notochord by exhibiting a markedly lower Brachyury1 level and lacking Fgf8/17/18 and Mnx expression. Despite slightly reduced levels, they maintain Chordin and Goosecoid expression. We designated this population as the PrCP and propose its homology with the vertebrate PrCP.

Recently, top-ranking DEG specifying the PrCP and notochord were identified from scRNA-seq data in zebrafish (*14*). We found orthologs of the corresponding genes in amphioxus and examined their expression in our dataset (Fig.2*A-D;* fig.S2A-B5 and **S2F1-F2).** At G4, PrCP-specific genes were predominantly expressed in the preaxial and axial mesendoderm (Fig.2*A*). By N0, most of the genes showed preferential expression in either the PrCP or notochord, reflecting patterns in zebrafish (Fig.2*A-B*). These observations suggest that the separation of the PrCP and notochord in amphioxus occurs during gastrulation, similar to vertebrates (*6, 9, 14*). Calculated cell state transition probabilities further support this notion (fig.S5*A*).

**Fig.2:**
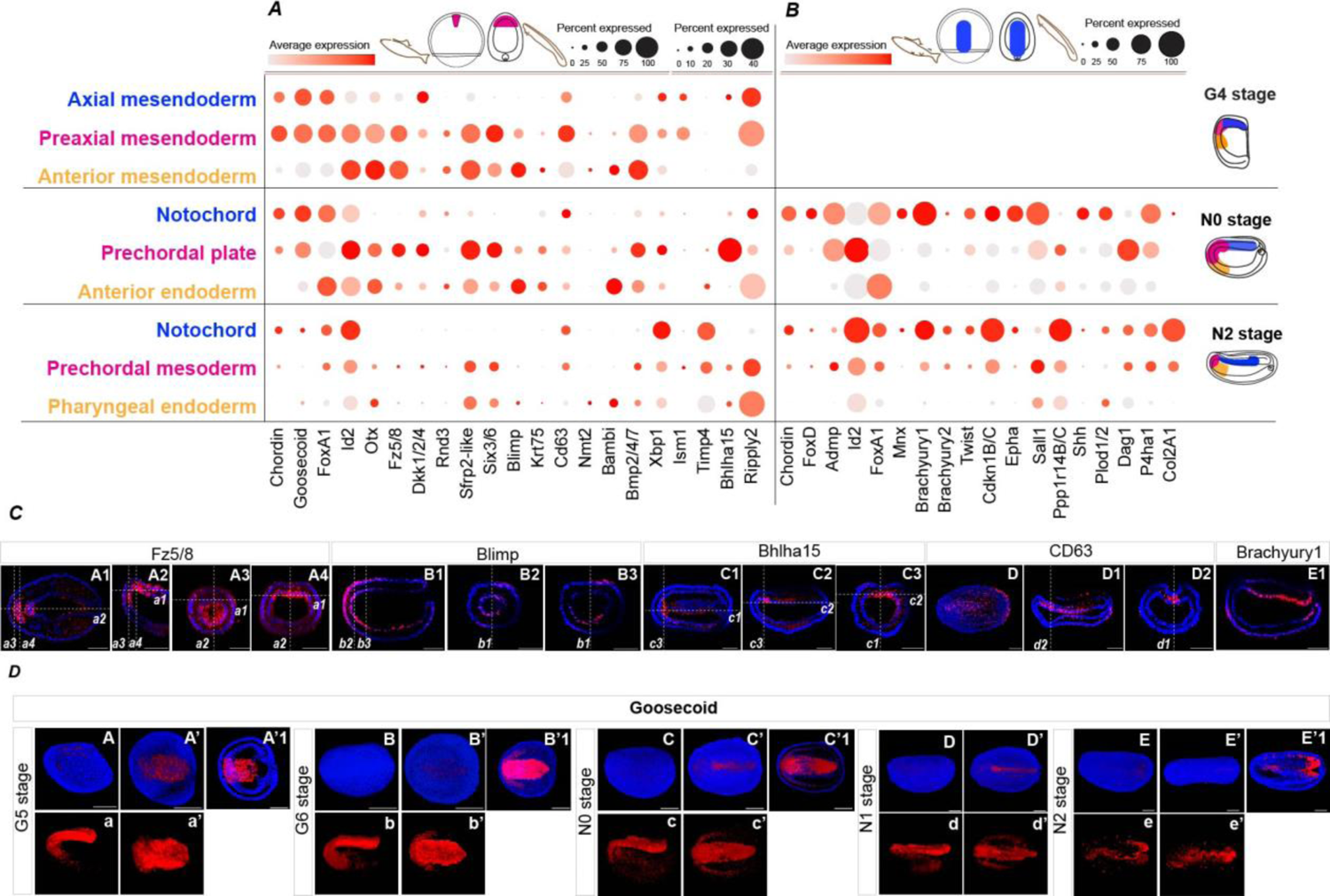
Single-cell analysis of PrCP and notochord markers in amphioxus, initially Identified in zebrafish, across the G4, N0, and N2 stages. The DotPlots depict the expression of PrCP (*A*) and notochord (*B*) marker genes for the selected cell populations. Schematic drawings of the amphioxus embryo in (*A*) and (*B*) illustrates the spatial position of individual clusters, colored in the same way as the names of the clusters. (*C*) Spatial expression of individual genes. (*C*A1-E1) Individual z-slices from z-stacks are indicated with numbered dashed lines (*a1-d2*). (*D*) Spatial expression of Goosecoid in the timeline from G5 to N2 stage. Lateral (*D*A-a, *D*B-b, *D*C-c, *D*D-d, *D*E-e) and dorsal (*D*A’-a’, *D*B’-b’, *C*C’-c’, *D*D’-d’, *D*E’-e’) 3D views of complete z-stacks. (A-A’, B-B’, D-D’, E-E’) display the signal with DAPI and (a-a’, b-b’, c-c’, d-d’, e-e’) display the signal alone. The section through (A’, B’, C’, E’) are shown in (A’1, B’1, C’1, E’1).

To understand the molecular differences between the notochord and PrCP at the N0 stage, we analyzed top DEG (fig.S6). The PrCP primarily exhibited genes linked to migration and cell motility, similar to the migrating PrCP cells in vertebrates (*8, 34*). The notochord displayed a distinct molecular signature, with a preponderance of genes involved in autophagy and energy metabolism.

At N2, we identified a cluster corresponding to the anterior dorsal mesendoderm, termed ‘prechordal mesoderm’ (Fig.1*N2*A-B). This population is marked by high Six3/6, Dmbx, and Bmp2/4 expression, but shows notably lower levels of Brachyury1 and Mnx compared to the notochord.

In our comparative analysis of vertebrate PrCP and notochord markers within amphioxus embryos, we observed a more distinct separation between the notochord and PrCP at the N0 stage compared to the notochord and prechordal mesoderm at N2 (Fig.2*A-B*). Certain notochord-specific markers, absent at N0, were present in the prechordal mesoderm at N2, though at lower levels (Fig.2*B*). The preaxial mesendoderm at G4 displays a transcriptomic profile more similar to vertebrate PrCP than the prechordal mesoderm at N2 (Fig.2*A*). Goosecoid persists in the PrCP until N1 but disappears by N2 (Fig.2*D*). These findings suggest that the amphioxus PrCP most closely mirrors its vertebrate counterpart during late gastrula to early neurula stages.

### Neural crest-like cells are present in amphioxus

At the N5 stage, a cluster termed ‘MNCC-like’ (Migratory Neural Crest Cells-like) was identified (Fig.1*N5*A-B), marked by high Elav, Islet, and Six3/6 expression. The abundant Six3/6 suggests that these cells are located in the embryo’s anterior domain (fig.S4J-J3). UMAP clustering revealed proximity between this cluster and the ‘PMNCC-like’ (Pre-migratory Neural Crest Cells-like) anterior neural tube population, characterized by increased Otx expression (Fig.3*A, E*; fig.S7*A*D1-D3). Computations of cell fate transition probabilities, which critically rely on transcriptomic profile similarity, indicate a strong transcriptional resemblance of MNCC to the anterior neural plate at N2 and PMNCC at N5. These computations also suggest that the most probable origin of MNCC lies within these domains (fig.S5,S8).

**Fig.3:**
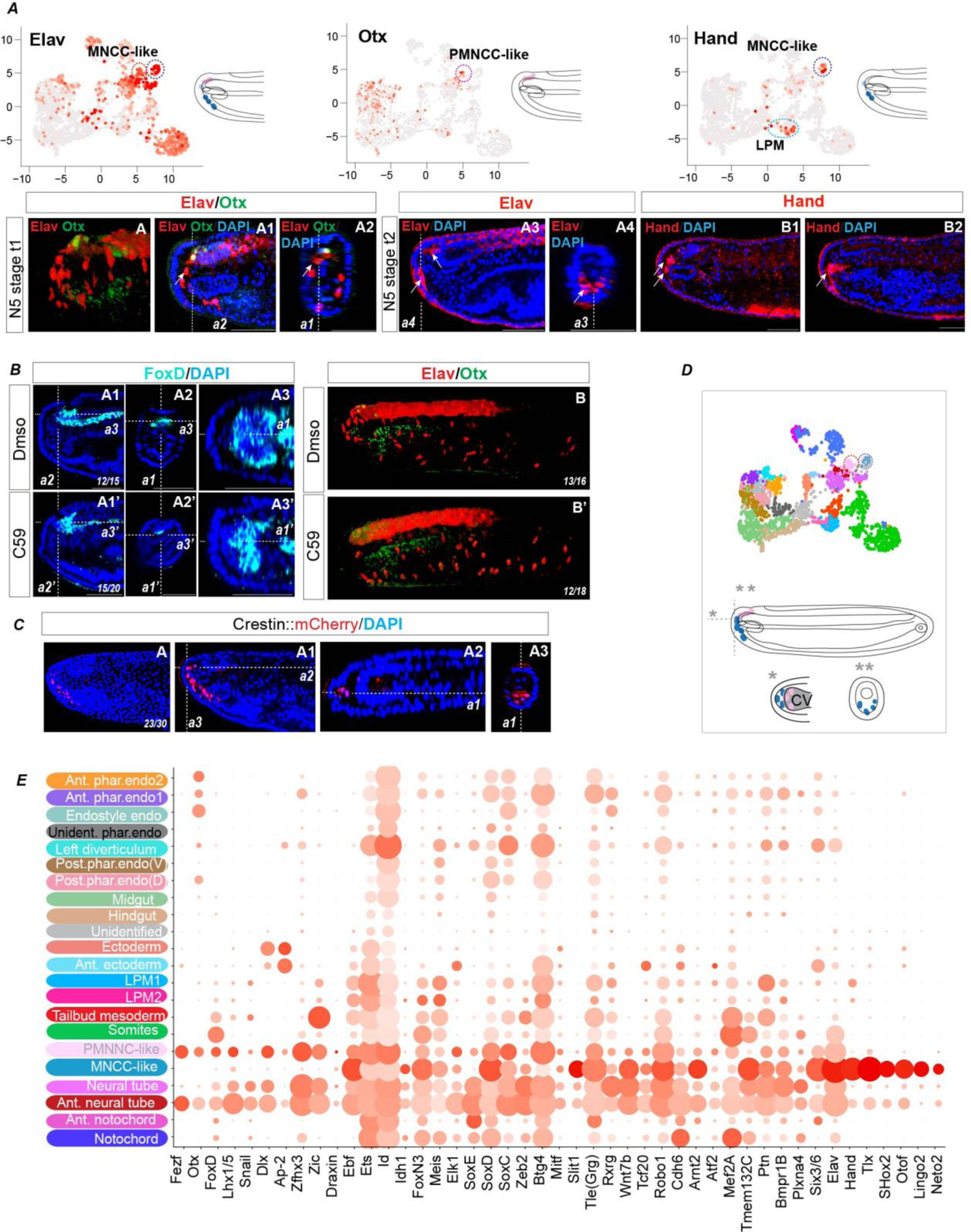
Neural crest-like cell are present in amphioxus. (***A***) The expression of Elav, Otx and Hand in PMNCC and MNCC on the UMAP plot and in the context of the embryo at N5. Arrows in (**A-B2**) indicate the expression in MNCC. (***B***) Wnt signaling inhibition blocks the emergence of Elav+ MNCC and FoxD+ PMNCC. (***C***) The activity of Crestin::mCherry in the embryos at N5. (***A*A**, ***B*B-B’**, ***C*A**) Lateral views of 3D-recontstruction of the complete z-stack. The position of individual stacks (***A*A1-B2**, ***B*A1-A3’**, ***C*A1-A2**) are indicated with dashed lines, marked as ***a1-a4***. (***D***) The dashed circles on the UMAP plot highlights PMNCC-like and MNCC-like clusters, with schematic drawings illustrating its spatial position in the amphioxus embryo. CV – cerebral vesicle. (***E***) The DotPlot illustrates DEG within PMNCC and MNCC.

ScRNA-seq analysis and double immunostaining revealed Elav co-expression with Otx in PMNCC, and a distinct strong expression of Elav alone in cells between the ectoderm and endoderm (Fig.3*A*A-A2). We observed dynamic changes in Elav expression at N5, particularly a shift in the concentration of Elav+ cells from the dorsorostral to the rostral region (Fig. 3*A*A3-A5).

MNCC also exhibited heightened expression of Hand (Fig.3*A*B1-B2). In vertebrates, Hand is associated with the cranial NC and lateral plate mesoderm(LPM) (*29, 35*), a pattern mirrored in amphioxus where Hand is notably expressed in the LPM (*36*) (Fig.3***A***). Additionally, Hand expression was especially strong in the MNCC-like cluster, the only population in our scRNA-seq dataset exhibiting extensive co-expression of Hand and Elav.

Analysis of DEG in MNCC identified genes including Shox2, Tlx, Prdm12, Trk, Scrt2, Skor2, Pou4f3, Neto2, Lingo2, Otof, and Runx1 (Fig.3*E*; fig.S7*E*). These markers in vertebrates typically signify developing sensory neurons within sensory ganglia (*27, 37–40*). Notably, vertebrate sensory ganglia originate from NC and cranial placodes.

Further examination of DEG in MNCC revealed numerous orthologs to genes associated with developing or migrating NC, including Ebf, Meis, FoxN3, Slit1, Robo1, Ets, Id, Idh1, Zeb2, Elk, Mitf, Rxrg, Itgb1, Ptn, Tle, Tmem132C, Bmpr1B, Arnt2, Rho and Adam genes (*24, 28, 30, 32, 41, 42*) (Fig.3*E*; fig.S7*A*D1-E9,*E*). While SoxE is absent, SoxD and SoxC are present, with SoxD being highly elevated. In vertebrates, the representatives of SoxC and SoxD groups are expressed in NC progenitors or derivatives (*43, 44*). The PMNCC-like population, likely a precursor to MNCC, exhibits a similar gene profile and additional vertebrate NC markers like FoxD, Lhx1, Ets, Snail, Zic, Btg, Dlx, Ap-2, Atf2, and Zfhx3 (Fig.3*E*; fig.S7*A,E*). Previous studies have shown that from the early neurula stage, amphioxus has Elav+ cells in the ventral ectoderm, which migrate between the ectoderm and endoderm (*45, 46*). In our scRNA-seq data, Elav+ cells were identified in the ectoderm at the G4, N0 and N5 stages. UMAP clustering and the calculation of cell fate transition probabilities revealed that these cells are distinct from the MNCC-like population (Fig.1; fig.S8).

Wnt signaling promotes NC induction and affects pre-placodal ectoderm induction in vertebrates (*32*). In our study, we noted high expression of Wnt7b and Tcf20 in PMNCC and MNCC-like populations, with Lrp6 and Dkk1/2/4 also elevated in PMNCC (Fig.3*E*; fig.S7*E*). Dkk1/2/4, orthologous to vertebrate Dkk1 and Dkk2, have contrasting roles in NC development: Dkk2 activates Wnt/β-catenin signaling for NC specification (*47*), while Dkk1, when secreted from the PrCP, inhibits it in the rostral neural plate border (*48*).

To explore the effect of Wnt signaling on PMNCC and MNCC, we applied the pathway inhibitor C59 from the N1 stage. Low concentrations of C59 did not noticeably alter embryo morphology but led to reduced FoxD expression in PMNCC and decreased Elav in both MNCC and PMNCC, while other Elav-+ cells remained unaffected (Fig.3*B*). Higher doses disrupted the overall development, but ectodermal cells persisted (fig.S7*D*). These findings suggest that PMNCC and MNCC populations are differently regulated by Wnt signaling compared to ventral ectoderm sensory cells.

To further examine our hypothesis that amphioxus MNCC-like population resembles the vertebrate NC, we utilized a trans-species transgenic reporter assay involving *crestin*, a zebrafish-specific gene expressed exclusively in NC (*49*). Previous studies have demonstrated that a regulatory element located 4.5 kb upstream of the predicted *crestin* open reading frame when linked to a reporter gene, accurately marks emerging and migrating NC (*50*). In transgenic amphioxus embryos, Crestin::mCherry reporter gene activity was present in the MNCC-like cells at N5, but absent in the most anterior neural tube (Fig.3*C*). At the N3 stage, the reporter signal appeared only sporadically in individual cells of the anterior neural plate (fig.S7*C*). In zebrafish, the transcription factors Sox10, Ap-2 and Mitf/Myc are crucial for *crestin* activity in NC (*50*). In the MNCC, while Mitf was present, Ap-2 was only found in PMNCC. Despite the absence of SoxE, other family members like SoxD and SoxC might bind to the SoxE motif, particularly given SoxD’s flexible DNA binding sequence (*51*).

Together, our data suggest that the migratory population, which exhibits features of the migrating and differentiating NC, originates from the specific domain in the anterior neuroectoderm.

### PrCP development in amphioxus is regulated by Nodal and Hh signalling

Motivated by evidence of Nodal (*7, 9*) and Hh signalling (*11, 31*) critical roles in PrCP development in vertebrates, we performed a pharmacological inhibition of these pathways in amphioxus to explore conservation of the respective PrCP programs at the gene regulatory level.

We conducted pharmacological treatments at the gastrula stage to target the early separation phase of the PrCP from the axial mesendoderm, focusing on PrCP formation rather than organizer formation. After applying the Nodal inhibitor at G4, we observed no significant reduction in Six3/6 expression in the PrCP at N1, nor any morphological deficiencies in larvae (Fig.4*A*). In contrast, G2 stage treatment eliminated Six3/6, Goosecoid, and FoxA1 expression in the PrCP (Fig.4*A*A-g’), significantly shortened the notochord’s anterior tip in larvae, and resulted in a deficient cerebral vesicle (Fig.4*A*I-T’). Expressions of Six3/6 and Fz5/8 (Fig.4*A*L-P2) diminished in the notochord’s anterior tip and anterior cerebral vesicle, while Lhx3 was absent from the ventral domain of the cerebral vesicle (Fig.**4*A*R-T’**). The absence of Lhx3 expression in the left diverticulum in G4-treated embryos aligns with previously described disruptions in left-right asymmetry during neurulation (*52*).

**Fig.4:**
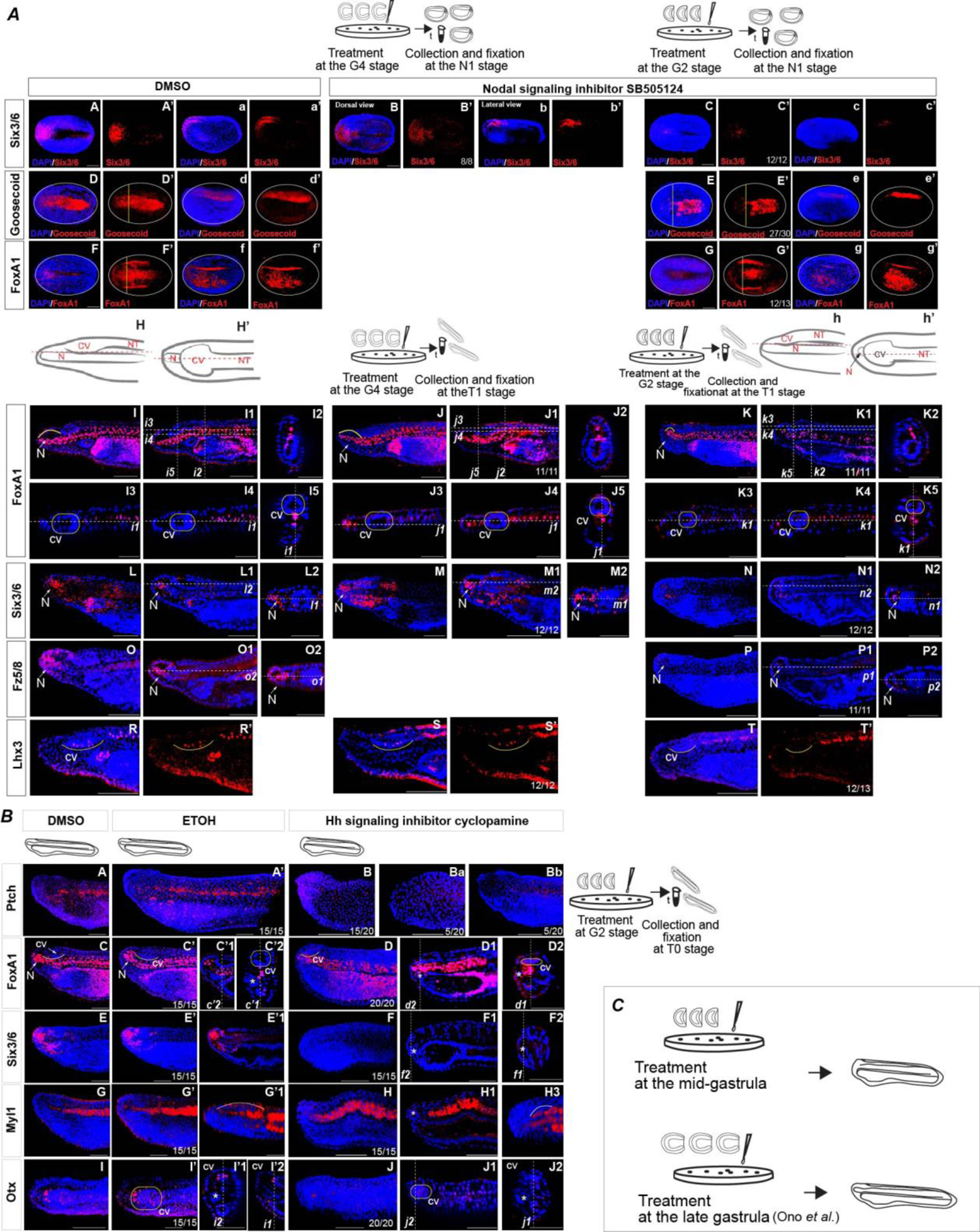
Inhibition of Nodal and Hedgehog (Hh) signaling pathways at the early gastrula stage impairs the development of rostral structures in a developing amphioxus. (*A*) Inhibiting Nodal signaling at G2, but not at G4, downregulated Six3/6, Goosecoid, and FoxA1 expression in PrcP by N1 (A-g’), along with a shortened anterior notochord (arrows) and an impaired cerebral vesicle in larvae (I-T’). (*B*) Cyclopamine treatment led to a shortened rostrum, downregulation of the Hh signaling target gene Ptch (*B*A-Bb), elimination of FoxA1 (*B*C-D2) and Six3/6 (*B*E-F2) expression in the anterior notochord’s, and reduced Myl1 (*B*G-H3) levels in the anterior somites. Six3/6 (*B*E-F2) and Otx (*B*I-J2) expressions were eliminated in the cerebral vesicle. (*C*) Schematic illustration depicting distinct effect of cyclopamine treatment at the mid-gastrula and late gastrula on head morphology. 3-D views of the complete z-stack of the entire embryo are presented in (*A*A-g’), while the head region is depicted in (*A*I, *A*J, *A*K, *A*L, *A*M, *A*N, *A*O, *A*P, *A*R-T’) and (*B*A-Bb, *B*C, *B*C’, *B*D, *B*E, *B*E’, *B*F, *B*G-H3, *B*I, *B*I’, *B*J). Asterisks in (*B*) indicate left diverticulum. The positions of individual z-slices (*A*I1-I5, *A*J1-J5, *A*K1-K5, *A*L1-L2, *A*M1-M2, *A*N1-N2, *A*O1-O2, *A*P1-P2) and (*B*C’1-C’2, *B*D1-D2, *B*E’1, *B*F1-F2, *B*I’1,-I’2, *B*J1-J2) from the complete z-stacks are indicated with dashed lines (*Ai1-i5, Aj1-j5, Ak1-k2, Al1-l2, Am1-m5, An1-n2, Ao1-o2, Ap1-p2*) and (*Bc’1-c’2, Bd1-d2, Bf1-f2, Bi’1-i’2, Bj1-j2*). CV – cerebral vesicle, NT – neural tube, N – notochord.

Administering the Shh signaling inhibitor cyclopamine at the G2 stage inhibited Ptch, a target of Hh signaling, and led to severe phenotypic malformations in larvae (Fig.**4*B***). These included the absence of the notochord’s anterior tip, defects in anterior neural tube closure, pharyngeal endoderm, and the first pair of somites. FoxA1 and Six3/6 expression vanished in the anterior notochord (Fig.**4*B*C-F2**), Myl1 was reduced in anterior somites (Fig.**4*B*G-H3**), and Otx was absent from the anterior cerebral vesicle (Fig.**4*B*I-J2**). The defects must be related to Shh activity in the preaxial mesendoderm at mid gastrula, as late gastrula cyclopamine application didn’t cause similar effect (*53*) (Fig.**5*C***). No severe rostral malformations were observed in amphioxus upon the Hh signalling components knockout, including Shh (*54*), Smo and Ptch (*55*), possibly due to the presence of maternal mRNAs or other Patched/Shh-like genes (fig.**S9**).

**Fig.5:**
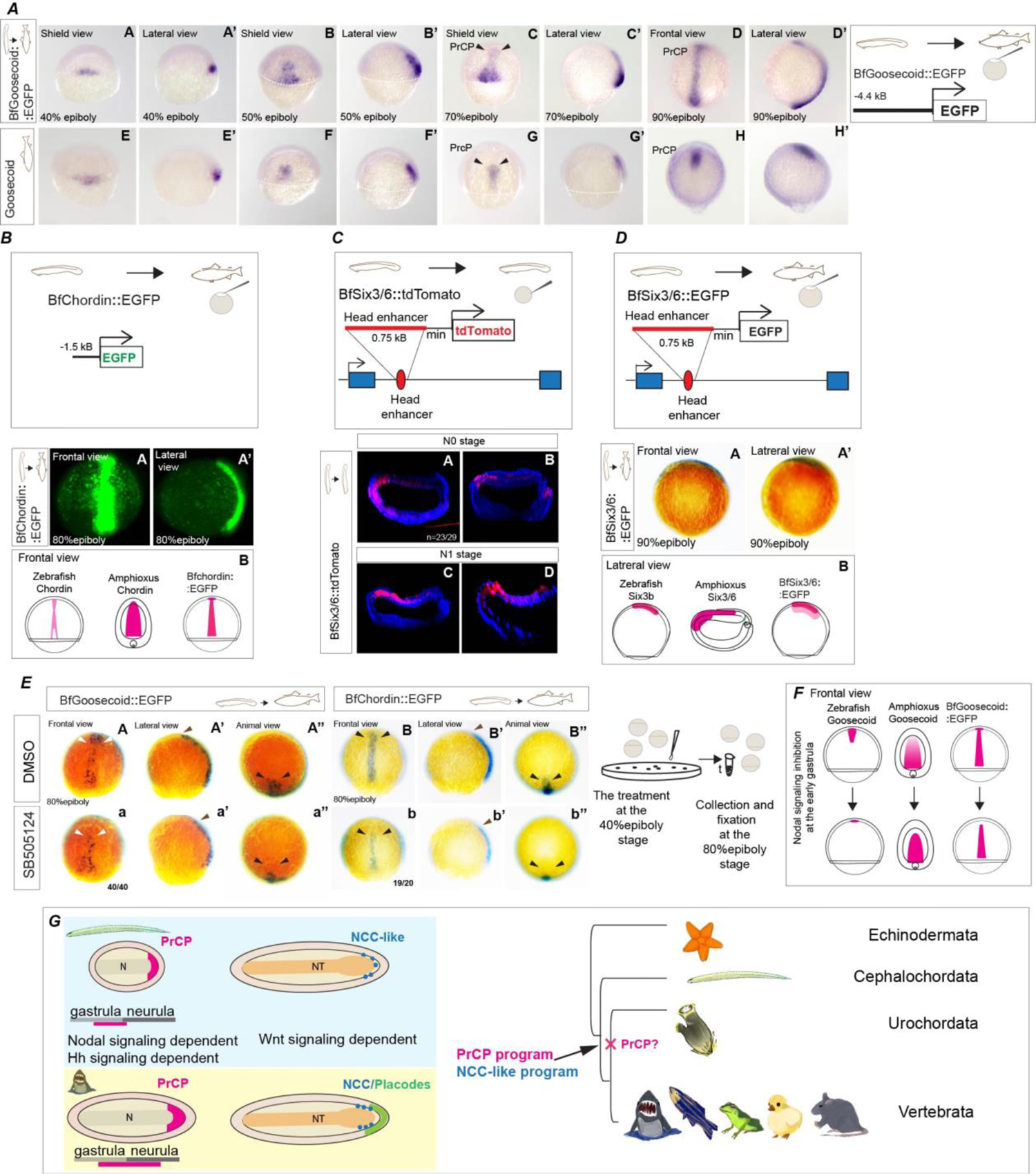
Functional conservation of prechordal plate development at the cis-regulatory level between amphioxus and zebrafish. (*A*A-D’) Activity of amphioxus Goosecoid cis-regulatory sequence in stable transgenic zebrafish embryos. (*A*E-H’) Expression of zebrafish Goosecoid. (*B*) Activity of amphioxus Chordin cis-regulatory sequence in stable transgenic zebrafish embryos. (*C-D*) Activity of amphioxus Six3/6 cis-regulatory sequence in transient transgenic amphioxus (*C*) and stable transgenic zebrafish embryos (*D*A-A’). (*B*B and *D*B) Schematic illustration comparing the activity of amphioxus Chordin and Six3/6 cis-regulatory sequences with the endogenous Chordin and Six3/b expression in zebrafish, as well as Chordin and Six3/6 in amphioxus. 3D-views of substacks from z-stacks in (*C*A-D). Z-slices of the whole embryo in (*C*A-C) and of the anterior region in (*C*D). (*E*) Inhibition of Nodal signaling with SB505124 at the early gastrula stage downregulates the activity of BfGoosecoid::EGFP (*E*A-a’’) and BfChordin::EGFP (*E*B-b’’) in the PrCP of zebrafish. (*F*) Schematic illustration depicts the changes in Goosecoid expression in both zebrafish and amphioxus, as well as the activity of the amphioxus cis-regulatory sequence in the context of a zebrafish embryo, upon Nodal signaling inhibition. (*G*) Emergance of PrCP and NCC in a common ancestor of all chordates. N – notochord. NT – neural tube.

### Cis-regulatory conservation of PrCP development in chordates

To further explore the conservation of PrCP development in chordates, we employed cross-species transgenic reporter assays (*35, 56, 57*). Specifically, we examined the activity of reporters driven by amphioxus Goosecoid (BfGoosecoid::EGFP), Chordin (BfChordin::EGFP), and Six3/6 (BfSix3/6::EGFP) cis-regulatory elements in developing zebrafish embryos.

BfGoosecoid::EGFP reporter gene activity was observed in the anterior mesendoderm and PrCP of amphioxus (fig.**S10*A***) and zebrafish embryos (Fig.**5*A***). This reporter also exhibited strong activity in the axial mesoderm, similar to its endogenous pattern. Likewise, BfChordin::EGFP (*18*) drove expression in the axial domain and PrCP at the 80% epiboly stage in zebrafish (Fig.**5*B*A-A’**), more closely aligning with the endogenous expression of Chordin in amphioxus than with that in zebrafish (*58*).

A recent functional annotation of the amphioxus genome using ATAC-seq identified potential cis-regulatory DNA elements (*56*). Testing three open chromatin regions in the amphioxus Six3/6 locus at gastrula to mid neurula stages (fig**.S10*C***), revealed an enhancer matching the endogenous anterior ectoderm and PrCP-specific expression of Six3/6 (Fig.**5*C*A-D**). In zebrafish embryos, this enhancer was active in the PrCP and anterior ectoderm domain (Fig.5DA-A’), resembling the expression pattern of endogenous zebrafish Six3b (*59*) (Fig.**5*D*B**). Treating zebrafish embryos with a Nodal signaling inhibitor at the early gastrula stage abolished BfGoosecoid::EGFP and BfChordin::EGFP expression in the PrCP, but left axial mesoderm expression unaffected (Fig.**5*E***, fig.**S10*B***). This response mirrors amphioxus results under similar conditions (Fig.**4*A***) and aligns with zebrafish patterns, where endogenous Goosecoid is lost from the PrCP after the treatment the at the early gastrula-stage (*7*) (Fig.**5*F***). These results provide additional support for the existence of an evolutionarily conserved regulatory program underlying PrCP development in chordates (Fig.**5*G***).

## Discussion

### Prechordal plate as an ancestor feature of chordates

Our study contributes to the debate on vertebrate head evolution by substantiating the ancestral origin of the PrCP in chordates. The obtained data suggest that the PrCP in amphioxus is specified and functions during the late gastrula to early neurula stages. By the N2 stage, the partitioning of the anterior neural plate in amphioxus resembles that in vertebrates (*21*), indicating that vital brain regionalization signaling events likely occur during earlier stages, aligning with the period when the PrCP’s transcriptomic profile closely mirrors that of vertebrates.

In vertebrates, Nodal signaling from the PrCP is essential for its development and for establishing the hypothalamus (*60*). In amphioxus, cerebral vesicle deficiencies, including the absence of Lhx3 in the ventral domain, were noted only with treatment at the G2 stage, not G4. This underscores the late gastrula/early neurula period as critical for Nodal signaling role in anterior neuroectoderm patterning, given that transcription inhibition by Nodal inhibitors can start within 30 to 45 minutes (*7*). At N0, Nodal signaling is active in the PrCP but not in the anterior neuroectoderm (fig.**S11*A*A-C**), indicating its role in patterning from the PrCP during this phase. By N2, nuclear phospho-Smad2 is not detected in the prechordal mesoderm (fig.**S11*A*D-E**). In early amphioxus neurula stages, Nodal signaling shifts to regulating left-right asymmetry (*52, 54*) (fig.**S11*B***). Conversely, in vertebrates, the PrCP-specific genes expression endures longer (*61*), with asymmetrical Pitx expression starting later compared to the progression of neurulation and forebrain development.

Similarly, Hh signalling during the late gastrula/early neurula phase is required for forming the head mesoderm, endoderm and cerebral vesicle in amphioxus, likely influenced by signals from the PrCP and preaxial mesendoderm, where Shh is abundant but absent in the neuroectoderm (Fig.**1*G4***). Post-N0 stage, Shh expression in the anterior mesendoderm significantly decreases (*53, 54*) (fig.**S11*B***). In vertebrates, Hh signaling from the prechordal mesoderm is crucial for head development, a function extending over a significant duration of neurulation (*10, 11*).

Meister *et al.* observed differences between the anterior tip and the central domain of the notochord during amphioxus neurulation, raising a question about the transient presence of vertebrate-like PrCP (*22*). Our findings corroborate the observation of notochord regionalization at the early mid-neurula and late neurula stages, revealing separate clusters for the notochord anterior domain at N2 and N5. However, the structure we described here as homologous to the vertebrate PrCP is distinct, and by the N2 stage, it no longer exists in a *bona fide* vertebrate-like manner. In addition, at the late gastrula/early neurula stages, PrCP includes additional lateral and anterior dorsal mesendoderm domains.

The PrCP in amphioxus, a transient cell population, likely transitions into various cell types. Specifically, our computation of cell fate transition probabilities suggests that the N0 PrCP may transition to the pharyngeal endoderm and prechordal mesoderm at N2, which then contributes to the anterior notochord and the Hatschek’s left diverticulum (HLD) (fig.**S5**). This aligns with the concept of the PrCP being part of the anterior mesendoderm, encompassing both dorsal and lateral regions. Accordingly, revisiting Goodrich’s hypothesis that HLD is of mesodermal origin and serves as a homolog of the vertebrates’ cranial premandibular coelom (*62*) is pertinent, especially considering that the premandibular coelom is also derived from the PrCP (*12*).

Previous studies have shown the complex gene expression patterns within HLD (*62, 63*). Our cell fate transition probability analysis supports this complexity, indicating a probable contribution from the anterior neural plate, likely due to high transcriptomic similarity and implying a sharing of gene regulatory network features. Post-metamorphosis, HLD fuses with the surface ectoderm to form Hatschek’s pit, potentially analogous to the vertebrate adenohypophysis (*1, 64*), originating from cranial placode and endoderm (*1, 65*). This high transcriptomic similarity aligns with the hypothesis that certain transcription factors in the chordate lineage may have expanded from the endomesodermal regions of HLD to adjacent ectoderm (*1*).

### Identification of neural crest-like cells in amphioxus

Studies suggest the presence of sensory cells migrating from the ventral ectoderm (*45, 46*), with a significant fraction of their genes found in vertebrates’ dorsally forming sensory cells (*66*). The MNCC-like population closely resembles vertebrates’ placode/NC-derived sensory neurons and migrating NC. However, it is distinct from the ventral ectoderm-derived peripheral sensory cells and most likely originates from the anterior neural tube. Alternatively, MNCC may originate from the rostral or ventral anterior ectoderm. Our findings reveal that, unlike MNCC, Wnt signaling is not necessary for Elav expression in late neurula stage ventral sensory cells. A recent study indicated that similar treatment marginally increases Elav+ cells in the mid-neurula ventral ectoderm (*66*). Conversely, activating canonical Wnt signaling inhibits Elav and Tlx in the anterior ventral ectoderm (*66*). This indicates that the sensory cells’ development in the ventral anterior ectoderm follows a regulatory mechanism distinct from that in MNCC. Moreover, the reduced FoxD and Elav expression in PMNCC due to Wnt inhibition indicates the necessity of Wnt signaling for the specification of these cells, similar to MNCC. Notably, Elav, absent in the rostral domain during neurula stages, is detected in the rostral ectoderm in larvae (*17, 67*), alongside Islet (*36*). These Elav+ cells may correspond to the rostral sensory cells described by Lacalli et al.(*68*), potentially originating from the MNCC.

We also considered the possibility that MNCC originate from the dorsorostral ectoderm, as they share some genes with this region, though less than with the PMNCC (Fig.**3*E***). Immunohistochemistry revealed dorsal/rostral Elav+ MNCC in embryos starting at early N5 stage, absent at N3 or N4 stages (fig.**S7*B***), suggesting their emergence occurs during late N4 or early N5. Yet, the appearance of a population resembling PMNCC more than the anterior ectoderm (fig.**S8**) during this period is improbable.

We propose that PMNCC specification occurs during the second phase of neurulation, with delamination at the end of neurulation. The PMNCC-like population in our dataset likely includes both pre-migratory and delaminating cells, starting to express sensory neuron markers. In this context, the MNCC-like population more closely resembles cranial placodes morphogenesis, where delaminating cells predominantly adopt a neuronal fate (*69*). A key argument against placodes’ existence in amphioxus is the lack of key markers like Six1/2, Eya, and Six4/5 in the dorsal non-neural ectoderm during late gastrula/early neurula stages (*63*). Supporting this, our scRNA-seq data found no Six1/2/Eya/Six4/5-positive clusters in the ectoderm at G4, N0 and N5, nor any expression in the anterior ectoderm at N2.

In *Ciona intestinalis*, sensory cells arise from proto-placodal or proto-NC regions, with potential interconversion following genetic disruptions (*70–72*). In amphioxus, PMNCC-like cells are situated in the most anterior neural tube, in close contact with the anterior ectoderm. The signals from the dorsal anterior ectoderm may contribute to the specification of PMNCC. In vertebrates, cranial NC is not specified at the rostral neural plate border, likely due to Wnt signaling inhibitors from the PrCP (*32, 48*). However, amphioxus PrCP resembles vertebrate PrCP only during the late gastrula/early neurula stages, and at N5, Wnt inhibitors Dkk1/2/4 and Dkk3 are absent in the anterior notochord of amphioxus (fig.**S7*E***).

Some genes identified in amphioxus MNCC are not present in vertebrate sensory cells originating from NC or placodes. Notably, vertebrate Mitf is expressed in NC-derived melanocytes (*73*) and in melanocytes derived from Schwann cell progenitors (*74*). Likewise, Hand2 is expressed in various NC-derived cells, including sympathetic ganglion and enteric neurons (*29*). This suggests two possibilities: either amphioxus and vertebrates have different sensory cell specification programs, or amphioxus MNCC may give rise to more than just sensory cells.

In summary, we propose that some cornerstone innovations in vertebrate head development, such as the PrCP and NC, can be traced back to ancestral features within the chordate lineage (Fig.**5*G***).

## Acknowledgments

We are grateful to Halyna Klymets for technical support during the manuscript improvement, Lucie Kozmikova for her drawings of animal representatives and Veronika Noskova for amphioxus facility maintenance. We acknowledge the Light Microscopy Core Facility, IMG, Prague, Czech Republic, for their support with the confocal imaging presented herein.

## Author contributions

Conceptualization: IK

Methodology: IK, AM, ZK, ZK-Jr, SMM, JP, JK

Investigation: IK, AM, AZ, ZK, ZK-Jr, SMM, JP, JK, SM

Visualization: IK, SMM, JK

Funding acquisition: ZK

Project administration: ZK, IK

Supervision: IK

Writing – original draft: IK

Writing – review & editing: IK, ZK, JK

## Data and Code availability

The scRNAseq datasets generated in this study have been deposited in the Gene Expression Omnibus (GEO) database under accession code “PRJEB67341”. The detailed code applied for the analysis in this study is available as ZIP file from the corresponding author upon request. Upon acceptance of this manuscript, we will make the GitHub repository publicly available and provide the associated URL and DOI.

## Supplementary Materials

### Materials and Methods

#### Maintenance and amphioxus husbandry

Amphioxus *Branchiostoma floridae* adults were housed in 5 litres of natural seawater at a temperature of 28°C and were fed with algae on a daily basis. To induce spawning, the animals were transferred to a temperature of 18°C for at least 6 weeks before being exposed to a heat shock induced by elevating the temperature to 28°C for 24 hours. One hour before artificial sunset, the animals were separated into 0.5-liter plastic cups containing 30-50 ml of seawater. Most of the animals spawned within an hour after the lights were switched off. The resulting embryos were raised at a temperature of 25°C.

#### Oligonucleotides

Oligonucleotides used for the generation of constructs used in WMISH, protein expression, and transgenesis are shown in Table S1.

#### Antibody production

Amphioxus-specific antibodies were prepared as previously described(*75*). Selected protein coding sequences were cloned into pET42a(+) vector (Novagen) using primers shown in Supplementary materials **Table S1.** Proteins were expressed in BL21 (DE3) RIPL bacteria (Stratagene) and purified using Ni-NTA agarose beads (Qiagen). Three mice of the B10A-H2xBALB/CJ strain were immunized four times in 4 weeks’ intervals with 30 mg of purified protein in PBS mixed with Freund’s adjuvant (Sigma-Aldrich). The serum was collected 10 days after the 4th immunization.

#### Immunohistochemistry staining of amphioxus embryos

After fixation with 4% PFA/MOPS (0.1 M 3-(N-morpholino)propanesulfonic acid, 1 mM EGTA, 2 mM MgSO4, 0.5 M NaCl, pH = 7.5) for 15 minutes on ice, the embryos were transferred through a series of 30% and 70% methanol mixed with 1xTBS and 0.1% Triton-X100 (TBST) to 100% methanol and stored at −20°C. The immunostaining procedure followed the general protocol (*75*) with minor modifications, such as the use of 1xTBS buffer with 0.05% Triton in all washing solutions, a 10% BSA and 10% donkey serum blocking solution and donkey anti-mouse secondary antibodies (ThermoFisherScientific) at the dilution 1:500. The embryos were imaged with Leica SP8 or Dragonfly confocal microscope and processed with Fiji ImageJ analysis software.

*In-situ* hybridization of amphioxus and zebrafish embryos The methodology used in this study involved conducting *in-situ* hybridization of amphioxus embryos, as previously described (*76*). The embryos were imaged with Leica SP8 confocal microscope and processed with Fiji ImageJ analysis software.

All experiments with zebrafish were performed in compliance with the European Communities Council Directive of 24 November 1986 (86/609/EEC) and guidelines of the Institute of Molecular Genetics of the Czech Academy of Sciences approved by the Animal Care Committee (approval numbers 11/2018 and 16-2023-P).

Zebrafish embryos were fixed in 4% PFA/PBT solution overnight at 4°C while shaking, and then stored in methanol at −20°C. *In-situ* was performed as previously described(*77*). Coloured signal was developed by incubation in BM-Purple (Roche) or Vector® Blue Substrate Kit at room temperature (RT) and then stored in glycerol at 4°C. Embryos were imaged with Olympus SZX10 and data were processed with Fiji ImageJ analysis software.

#### Cloning of the reporter gene constructs, generation and analysis of stable lines in zebrafish

The regulatory sequences of amphioxus Goosecoid and Six3/6 genes were amplified from B.floridae genomic DNA with oligonucleotides shown in (Table S1). Enhancer sequences were inserted into pZED vector upstream of the zebrafish gata2a minimal promoter(*78*) and promoter sequences into promoter-less pZED derivative. Transgenesis was performed with Tol2 transposon/transposase method (*79*). A mixture of 30 ng/µl of transposase mRNA, 30 ng/µl of Tol2-based transgenic construct, and 0.05% phenol red was injected into the one-cell stage embryos. Two independent transgenic lines were generated for BfGoosecoid::EGFP and BfSix3/6::EGFP constructs. The embryos of F1 and later generations were analysed with *in-situ* hybridization. Previously generated transgenic line containing BfCordin::EGFP (*18*) was analysed with light sheet fluorescent microscopy. The embryos were imaged with a Zeiss Light Sheet Z.1 microscope and processed with Imarisx64 9.6.0 and Fiji ImageJ software.

#### Transient transgenesis in amphioxus

Intronic enhancer of amphioxus Six3/6 and zebrafish −4.5 kb *crestin* regulatory element were cloned upstream of the minimal actin promoter and tdTomato or mCherry reporter in the vector carrying *PiggyBac* transposon terminal repeats (*56*). The −4.4kb promoter of amphioxus Goosecoid was cloned into the promoter-less version of the same vector. For microinjection of amphioxus eggs, mixture of BfGoosecoi::tdTomato or Six3/6::tdTomato (200 ng/µl) or Crestin::mCherry with PiggyBac transposase mRNA (100 ng/µl) in 15% glycerol was used. Transgenic embryos were allowed to develop until G4, N0 or N1 stage, fixed overnight in 4% MOPS/PFA (pH = 8.0) at 4°C, stained with DAPI, mounted with Vectashield (Vector Laboratories), and analysed using Leica SP8 confocal microscope. The confocal images were processed with Fiji ImageJ.

#### Pharmacological treatment of zebrafish and amphioxus embryos

The Nodal signalling pathway was inhibited with SB505124. Zebrafish embryos were placed in E3 medium and allowed to develop until the 40% epiboly stage when SB505124 was added at a concentration of 25 µM. Subsequently, the embryos were fixed at either the 80% or 90% epiboly stage and analysed with in situ hybridization.

Amphioxus embryos were treated with 3µM SB505124 in sea water at either the G2 or G4 stage and fixed at the N1 or T1 stage for analysis. The expression of Six3/6, Goosecoid, and Fz5/8 genes was analysed with in situ hybridization, and FoxA1 expression was analysed with immunohistochemistry staining.

Hedgehog signalling pathway was inhibited with 4µM cyclopamine. The embryos were treated with DMSO, EtOH or cyclopamine at the G1 stage and allowed to develop until T0 stage for the analysis. The expression of Ptch, Six3/6, Myl1 was analysed with in situ hybridization, FoxA1 and Otx expression was analysed with immunohistochemistry staining.

Wnt signalling was inhibited with 3 and 10 µM C59. The embryos were treated at N1 and allowed to develop until N5 stage for the analysis. The expression of FoxD was analysed with in situ hybridization, Elav and Otx expression was analysed with immunohistochemistry staining.

#### Preparation of single-cell samples and 10x genomics single-cell library from amphioxus

For each sample, 15 to 20 morphologically normal embryos were randomly picked and transferred into 4 well plates pre-coated with BSA and filled with filtered sea water (FSW). The embryos were washed with Ca2+, Mg2+-free artificial sea water (Ca2+, Mg2+-free ASW, 0.5 M KCl, 2 M NaCl, 0.5 mM NaHCO3, 0.2 M Tris–HCl, pH=8, 0.33M Na2SO4). Then, the embryos were transferred into tubes pre-coated with 5% BSA in Ca2+, Mg2+-free artificial sea water and immediately dissociated with preheated 1 to 2.5% trypsin in Ca2+, Mg2+-free ASW for 3-10 minutes (from the gastrula to late neurula stages) at 37°C and 150 rcf, while pipetting to complete the dissociation. The digestion was then inhibited with 20% FBS.

Cells were collected by centrifugation at room temperature for 5 minutes at 100 rcf and then resuspended in ASW and filtered. The cell concentration and viability of each sample was checked using Trypan Blue solution (Sigma Aldrich) with the Automated Cell Counter TC20TM (Biorad). Samples with a viability above 90% were used for single-cell sequencing.

Samples containing the cells from the embryos at the G4, N0, N2 and N5 stage were collected and washed with 1.2 M glycine and then dissociated. Single-cell encapsulation, cDNA synthesis, and library preparation were performed using Chromium nextGEM single cells 3’reagent kit v3.1, samples were targeted to 5000 cells (Table S2).

Sequencing was performed on Illumina NextSeq500 with mRNA fragment read length of 130 bases. UMI (Unique Molecular Identifiers) were quality controlled and counted by Cell Ranger Single-Cell Software Suite 6.1.1 from 10 × Genomics using *B. floridae* genome (*80*) with added manually curated genome models from *B. lanceolatum* genome (https://genome.jgi.doe.gov/) and (GitHubrRpository > 10X_matrices > gene_id_conversion_table.csv).

#### Single-cell RNA-seq data analysis

Two single-cell RNA datasets on amphioxus were previously published, providing a context for our analysis. The first study (*81*), conducted on live cells, yielded a cell count similar to ours. The second, employing single-nucleus analysis from fixed cells, resulted in a cell count tenfold greater (*82*). Despite this substantial difference in cell numbers, our comparative evaluation of gene expression profiles showed minimal disparities (fig.S12). Importantly, in both our dataset and that of Ma *et al.* (*82*), we observed the co-expression of Six3/6 and Fz5/8 in the presumptive PrCP at the N0 stage, consistent with known biological patterns.

However, PrCP-specific genes like Dmbx1 and Bhlha15 were only found in scattered cells in the dataset of Ma *et al*. (*82*) and did not match their expected endogenous expression patterns. This omission suggests possible limitations in the annotation of the dataset.

More cells do not necessarily equate to a superior dataset or analysis. This is particularly significant in datasets using 10XGenomics technology, where around 5% of approximately 4000 cells are duplicates, presenting challenges for analysis.

In our study, despite having fewer cells, we successfully identified all relevant populations, later validating them through conventional in situ hybridization and immunohistochemistry techniques. Notably, in our N5 stage dataset, which contains the smallest number of cells, we achieved the highest depth coverage. This enabled us to reliably identify populations present in small numbers within the embryo, such as the left diverticulum or the anterior tip of the notochord.

At the N2 stage, we were unable to identify the ectoderm cell population. The typical ectoderm marker Ap-2 was only randomly expressed in 3 cells. In contrast, other analysed stages showed distinct and well-defined Ap-2 positive clusters. The challenges in identifying the N2 ectoderm could be attributed to the secretion of the hatching enzyme between the ectoderm and the fertilization envelope around this stage. During single-cell sample preparation, the ectodermal cells might have been excessively exposed to both the hatching enzyme and trypsin, leading to the digestion of cell membranes and compromising their condition. Considering that the sample demonstrated the viability of more than 90%, it is highly probable that ectodermal cells did not settle during centrifugation and were removed with the supernatant. As a result, we obtained a sample enriched with cells from the sub-ectodermal populations and were able to correctly identify and verify all non-ectodermal relevant populations within the N2 stage embryo.

Analysis of gene expression matrices, clustering, and projections construction were conducted using the Seurat package v4.3.0.1 in R. The datasets from the G4, N0, N2, and N5 stages were analysed individually, adhering to the parameters specified in (Supplementary information: Githubrepository > Individual_timepoints (local copy provided with submission)).

To identify cell populations, we utilized a semi-supervised approach for cluster identification and annotation. Initially, unbiased clustering was applied to delineate distinct clusters. Following this, the top differentially expressed genes within each cluster were assessed based on their spatial embryonic expression patterns, which were sourced either from previously published resources or our own investigations (figs.S1, S1a-d, S2, S2a-e, S3, S3a-e, S4, S4a-e). The cluster identities were ascertained through an analysis of the combinatorial spatial gene expression codes depicted in (Fig.1).

The head area was the main focus of our research. Therefore, extensive analysis of spatial gene expression was performed for the clusters corresponding to the anterior mesendoderm and preaxial mesendoderm at the G4 stage, anterior endoderm and prechordal plate at the N0 stage, prechordal mesoderm, pharyngeal endoderm, and anterior neural plate at the N2 stage, and anterior tip of notochord, left diverticulum, anterior neural tube, PMNCC and MNCC populations at the N5 stage. Extensive analysis of combinatorial spatial gene expression allowed us to directly annotate these clusters.

Due to the complexity of the datasets, some clusters were challenging to annotate accurately without further extensive spatial gene expression analysis. These clusters were not the primary focus of our research, and we did not conduct in-depth analysis on them. For instance, there were several clusters that corresponded to the subpopulation of somites or ectoderm, but their precise spatial localization was not determined. To streamline the analysis, we grouped some of these clusters together into a single ‘somites’ or ‘ectoderm’ cluster for simplicity (see GitHubRepository > Individual_timepoints)).

To infer probable fates of annotated cell types across timepoints, we utilized a modified URD approach used in (*14*). URD calculates cell fate transition probabilities by first utilizing defined timepoints, the initial and final stages in a dataset. It then critically employs the transcriptomic profiles of individual cells, analyzing changes in gene expression across these timepoints to determine the likelihood of cell fate transitions. First, we integrated expression data from individual stages using mutual nearest neighbors (MNN) algorithm (*83*). Then, diffusion map was generated based on the corrected data. The transition probabilities were then determined by the Euclidean distance between cells in gene expression space, as represented on a diffusion map, and converted to transition matrix. For subsequent pseudotime estimation within URD framework, we aimed to identify root cells (pseudotime=0) with more fidelity than simply selecting all cells from G4 stage. To achieve that, cell differentiation potential was computed using CytoTRACE algorithm(*84*) implemented in CellRank(*85*). As a root cells, we then identified G4 cells exhibiting CytoTRACE score < 0.15. As a next step, cell transition probabilities matrix was biased by the differences in each cell’s assigned pseudotime, favoring transitions that align with the direction of real time according to logistic function. Parameters of the logistic function were estimated based on overall pseudotime distribution in the dataset. Next, transition probabilities were averaged across groups of cells defined by cell types annotated during individual stages analyses and the stages of origin. Resulting transition matrix, representing probability of differentiation of individual cell types to cell types in consecutive stages, were z-scaled for more feasible interpretation and visualization (GitHubRepository > Transitions). To visualize the transition probabilities as shown in Fig. S5, we employed a series of data processing steps. First, transition matrix containing cell types of interest was converted to a directed graph with edge weights represented by scaled transition probabilities. To de-noise the graphs, we restricted edges to only connect to their respective consecutive stages (nodes). We further applied an arbitrary weight threshold—0.9 for the prechordal plate lineage and 1.5 for the crest lineage - while retaining the highest-weight edge to prevent nodes from becoming disconnected (GitHubRepository > Transitions).

**fig.S1:**
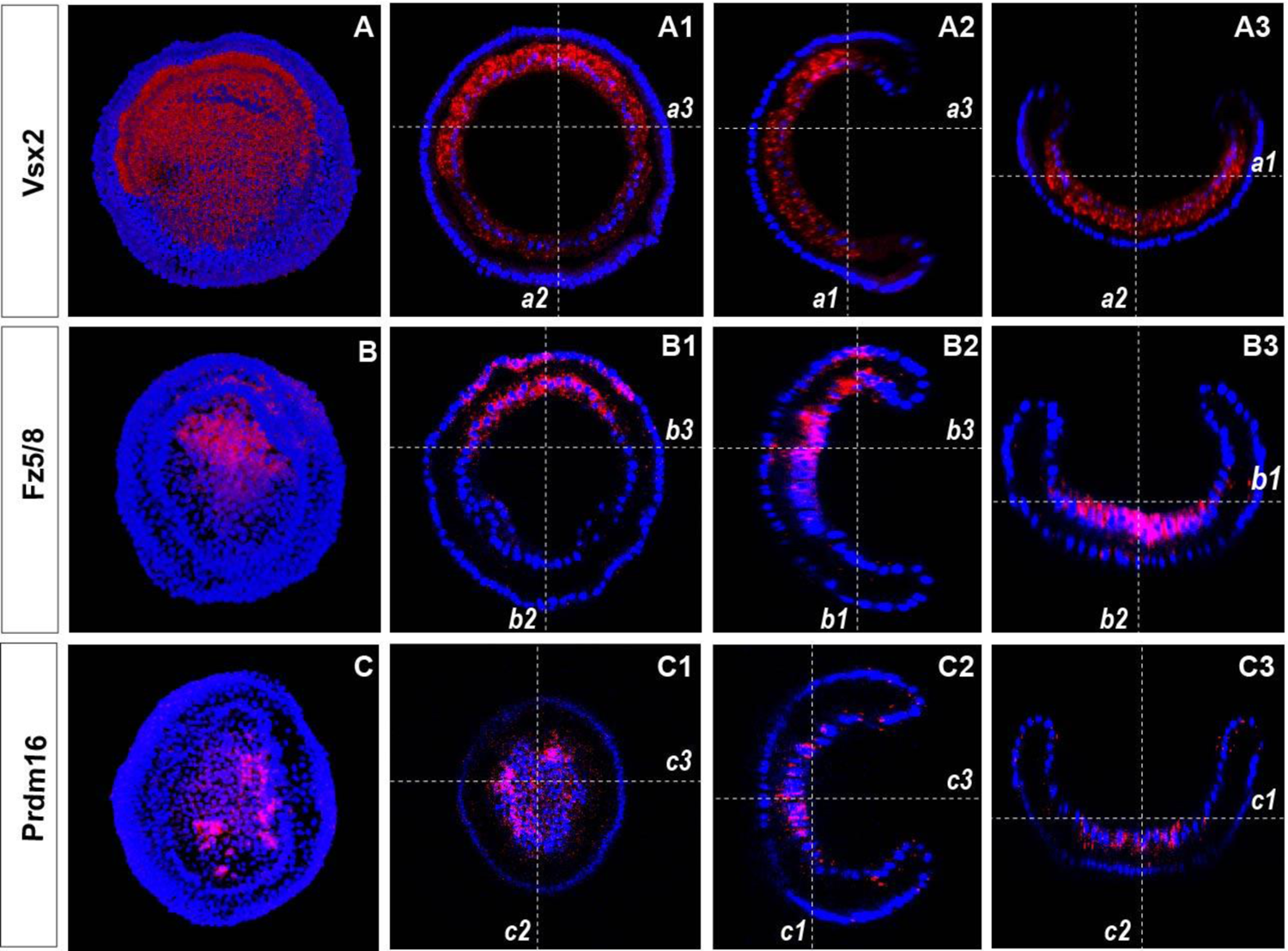
Spatial gene expression of selected genes at G4 stage. (**A**, **B**, **C**) Blastopore view of 3D-recontstruction of complete z-stack imaged with laser scanning confocal microscope Leica SP8. Individual z-slices from z-stacks are indicated with numbered dashed lines (***a1-a3, b1-b3, c1-c3***) and viewed in **A1-A3, B1-B3, C1-C3.**

**fig.S1a:**
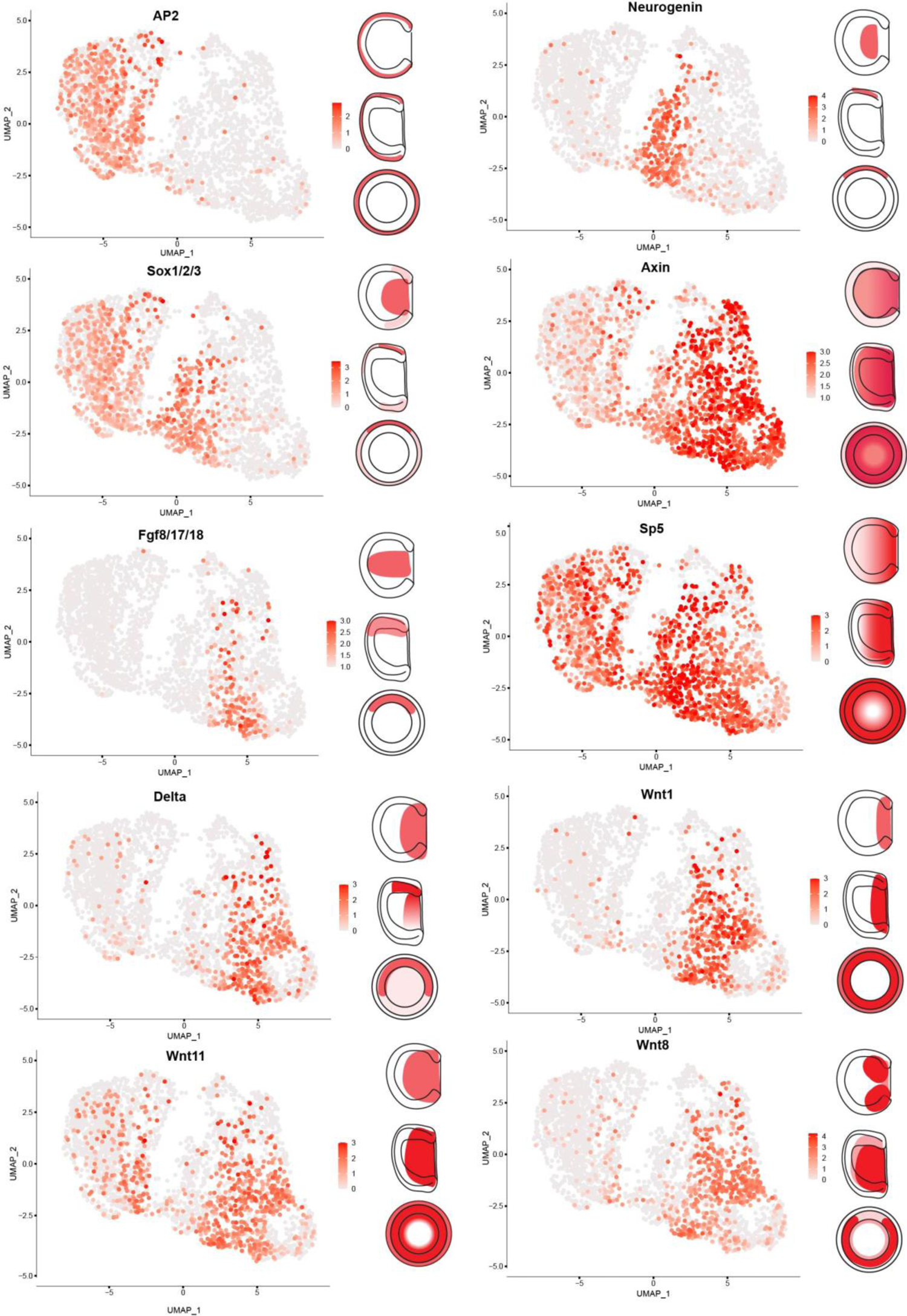
The feature plots showcase cells that are positive for each individual marker gene depicted in Fig.1G4, accompanied by schematic drawings illustrating the spatial gene expression from dorsal,

**fig.S1b:**
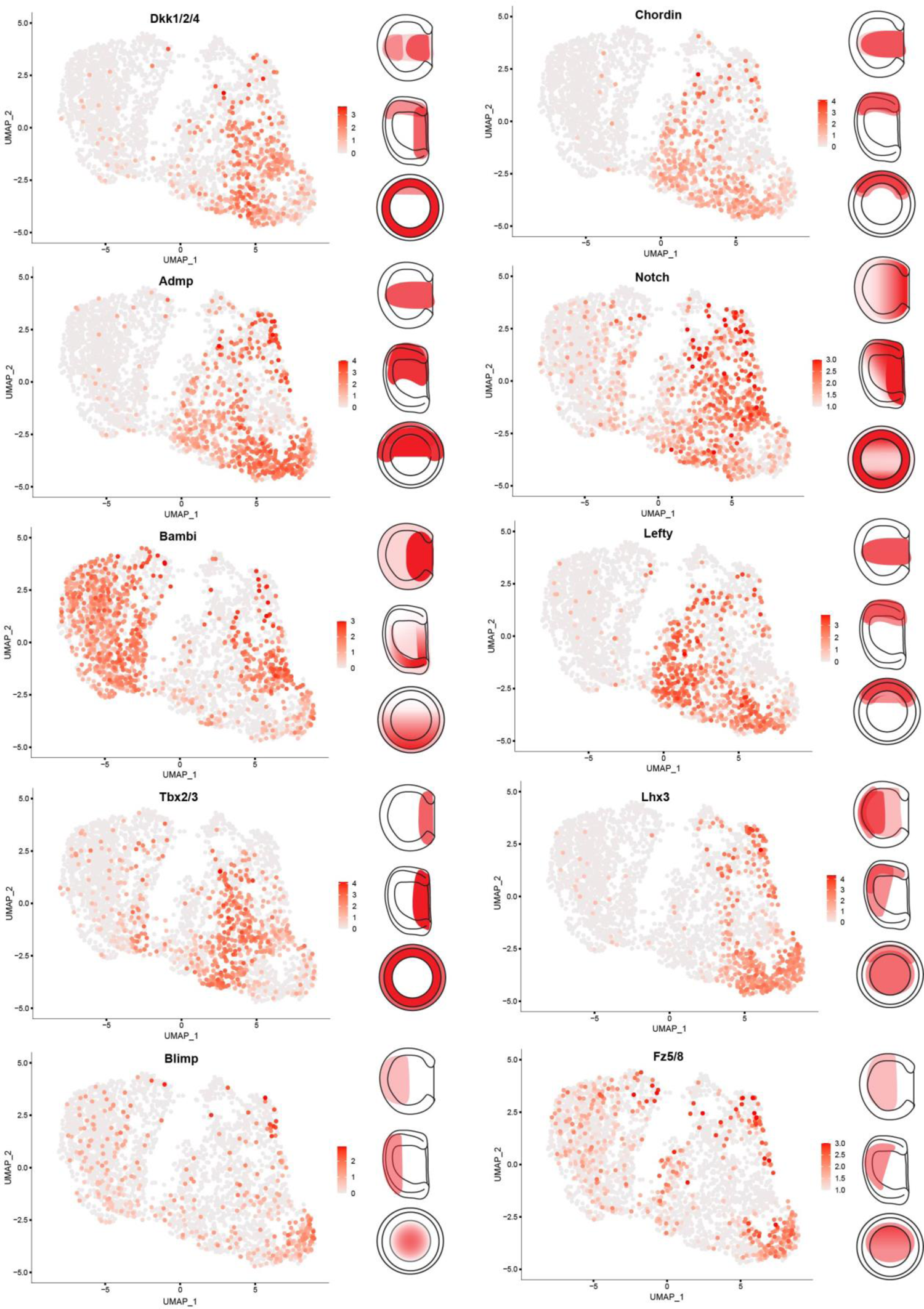
The feature plots showcase cells that are positive for each individual marker gene depicted in Fig.**1*G4***, accompanied by schematic drawings illustrating the spatial gene expression from dorsal,

**fig.S1c:**
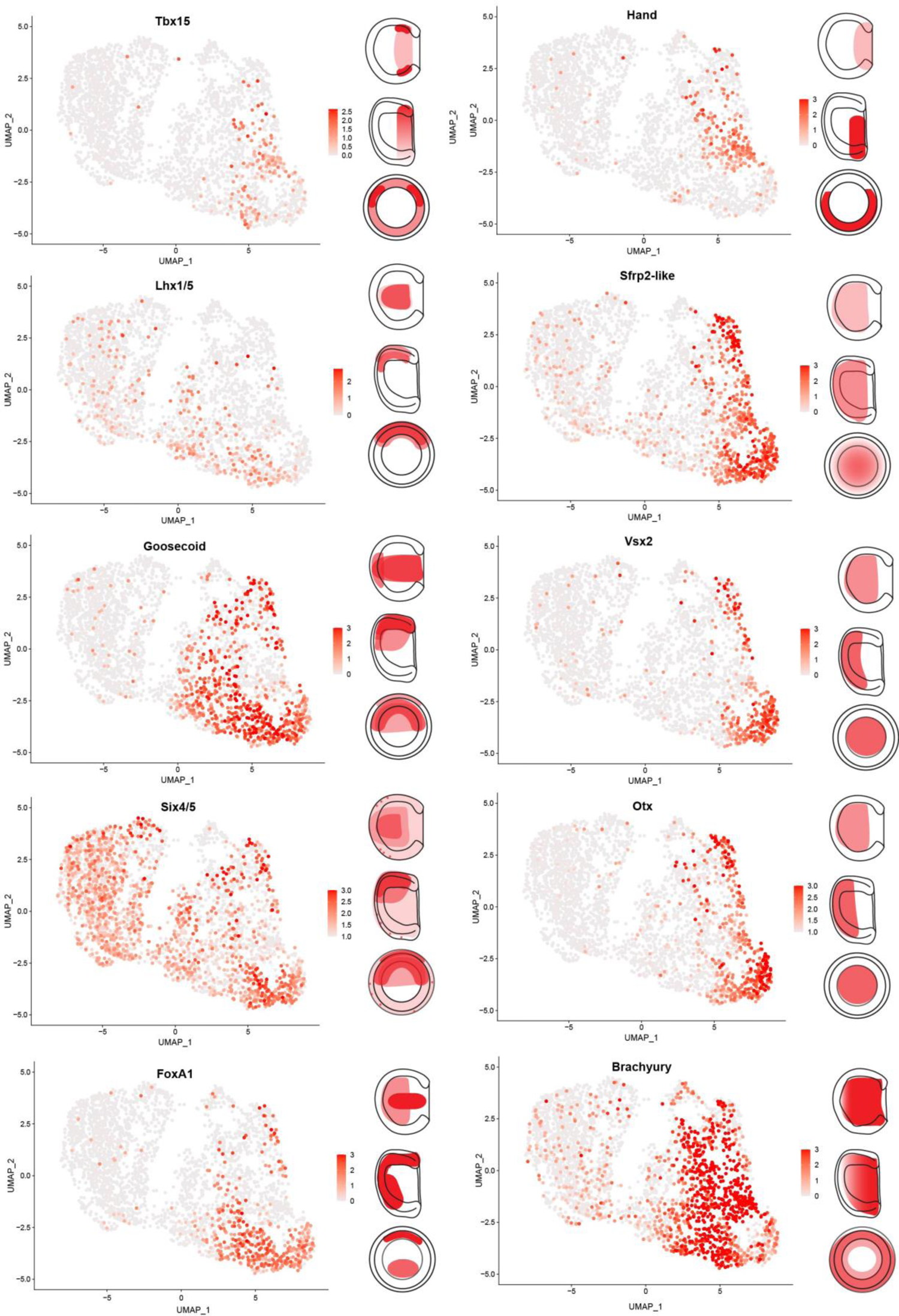
The feature plots showcase cells that are positive for each individual marker gene depicted in Fig.**1*G4***, accompanied by schematic drawings illustrating the spatial gene expression from dorsal,

**fig.S1d:**
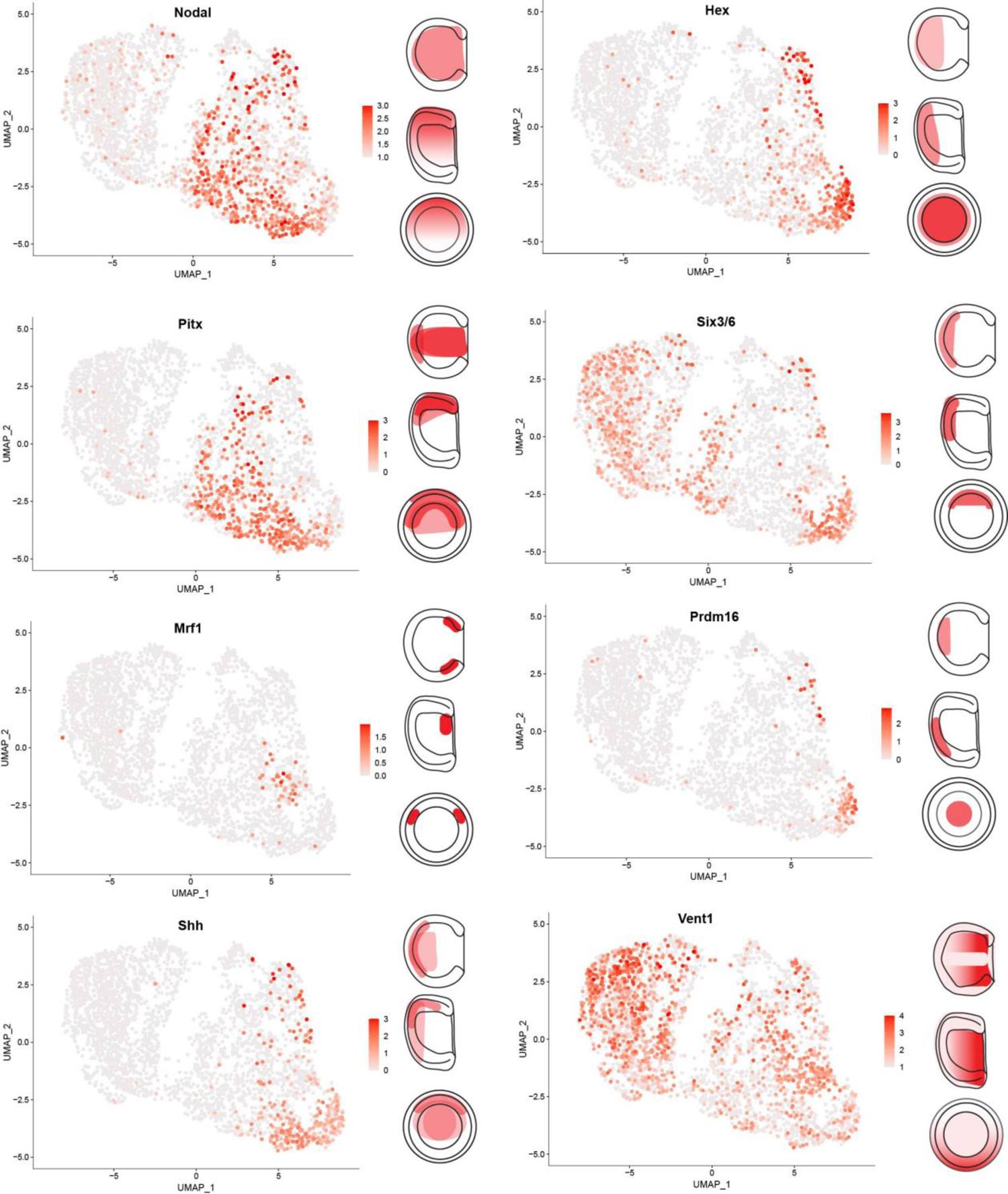
The feature plots showcase cells that are positive for each individual marker gene depicted in Fig.**1*G4***, accompanied by schematic drawings illustrating the spatial gene expression from dorsal, lateral, and blastopore views (from top to bottom).

**fig.S2:**
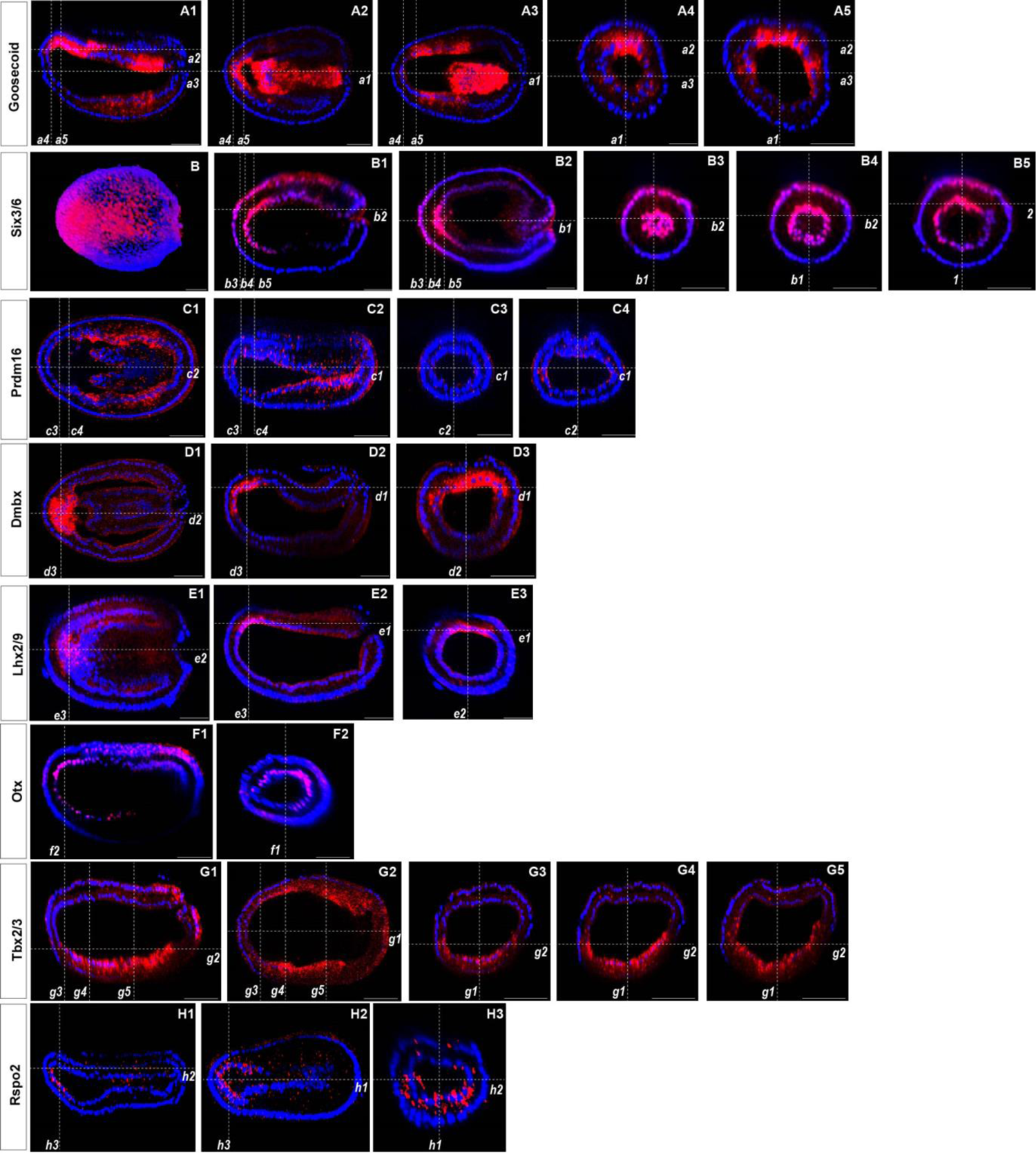
Spatial gene expression of individual genes at the N0 stage. The position of individual z-slices (**A1-H3**) from z-stacks imaged with laser scanning confocal microscope Leica SP8 are indicated with dashed lines (***a1-h3***). (**B**) Dorsal view of 3D-recontstruction of complete z-stack.

**fig.S2a:**
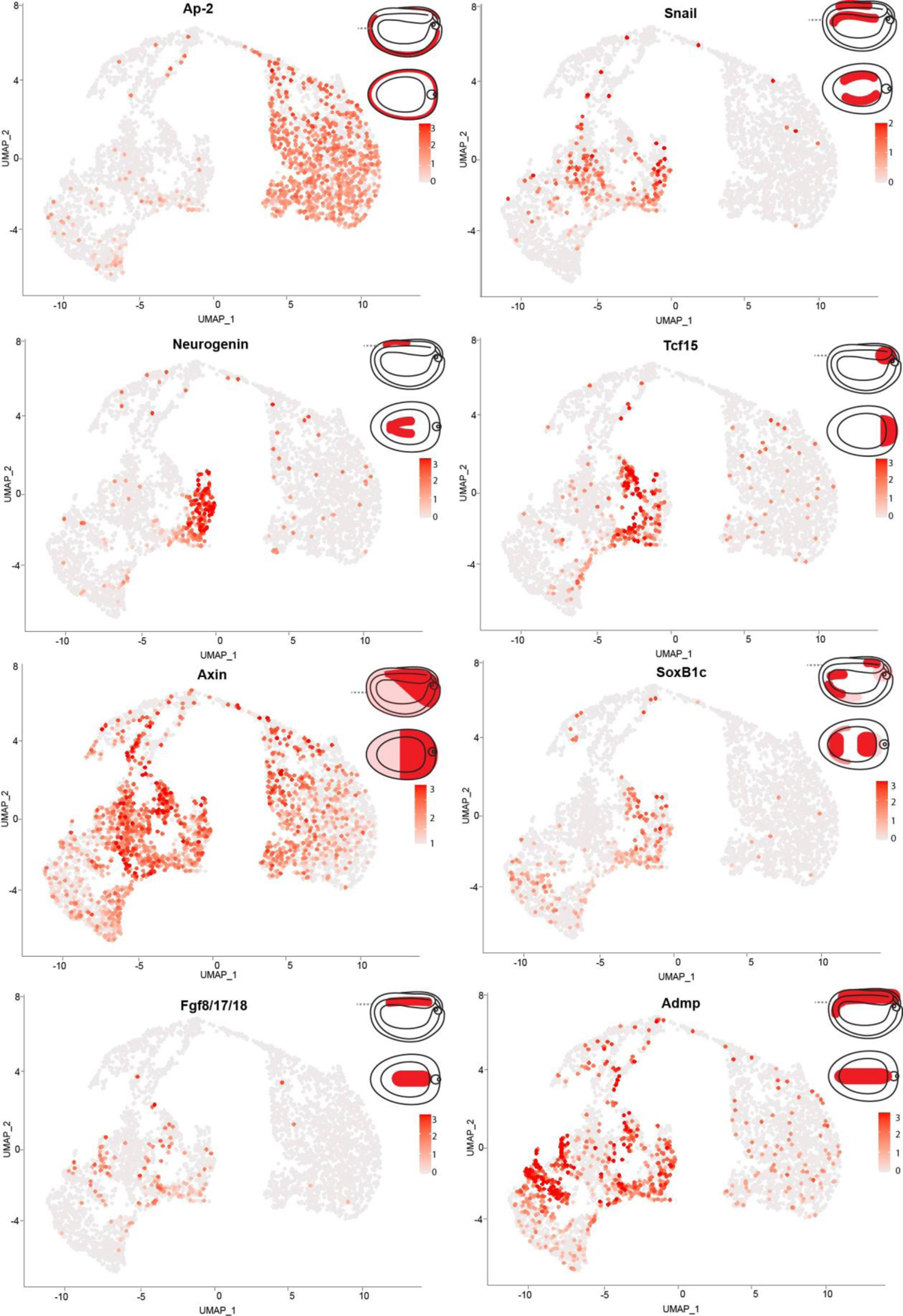
Feature plots of cells which are positive for each individual marker gene from Fig.**1*N0*** with schematic drawings of spatial gene expression (starting from the top: lateral view, dorsal view section trough the position marked with dashed line in lateral view).

**fig.S2b:**
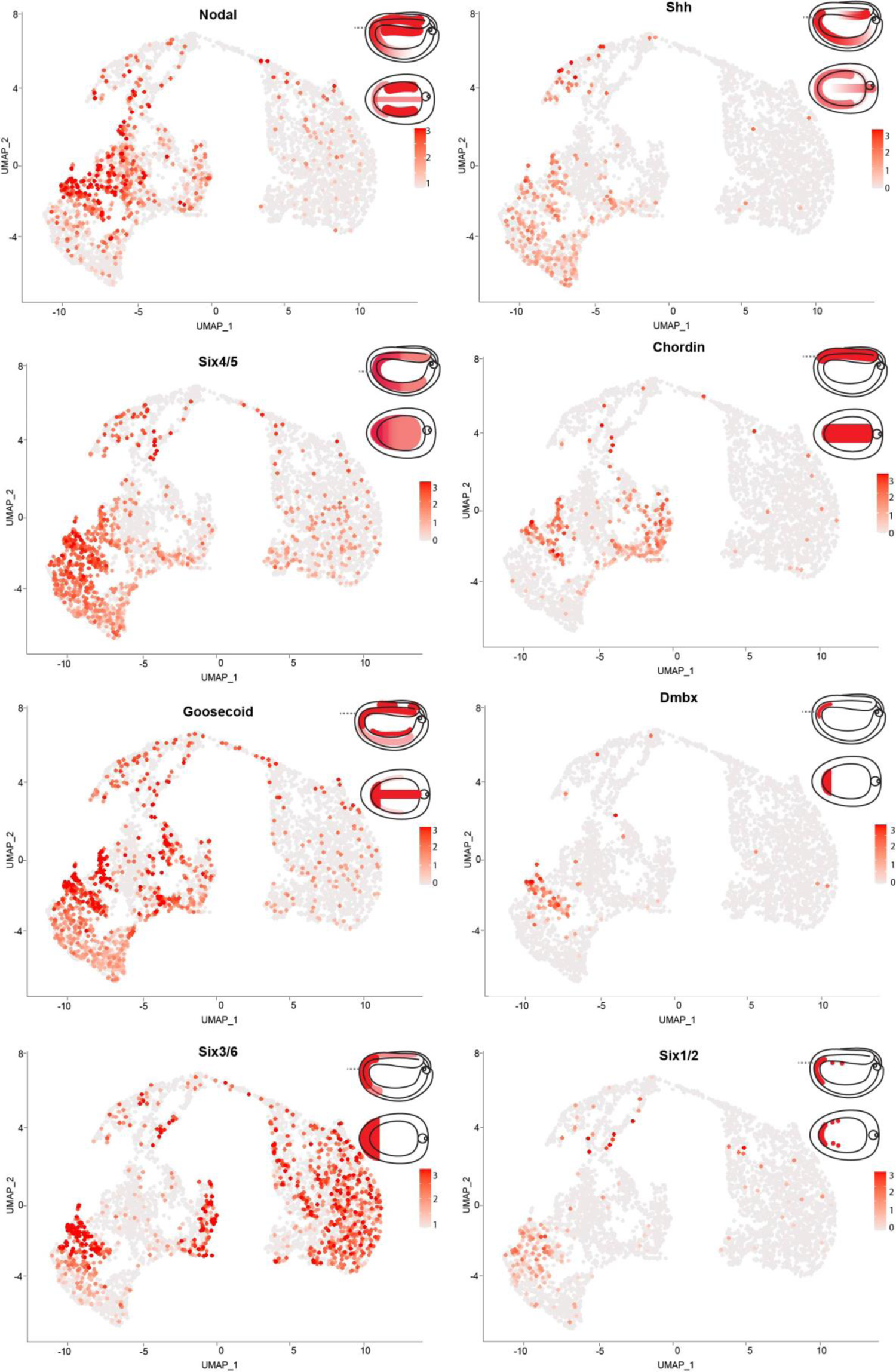
Feature plots of cells which are positive for each individual marker gene from Fig.**1*N0*** with schematic drawings of spatial gene expression (starting from the top: lateral view, dorsal view section trough the position marked with dashed line in lateral view).

**fig.S2c:**
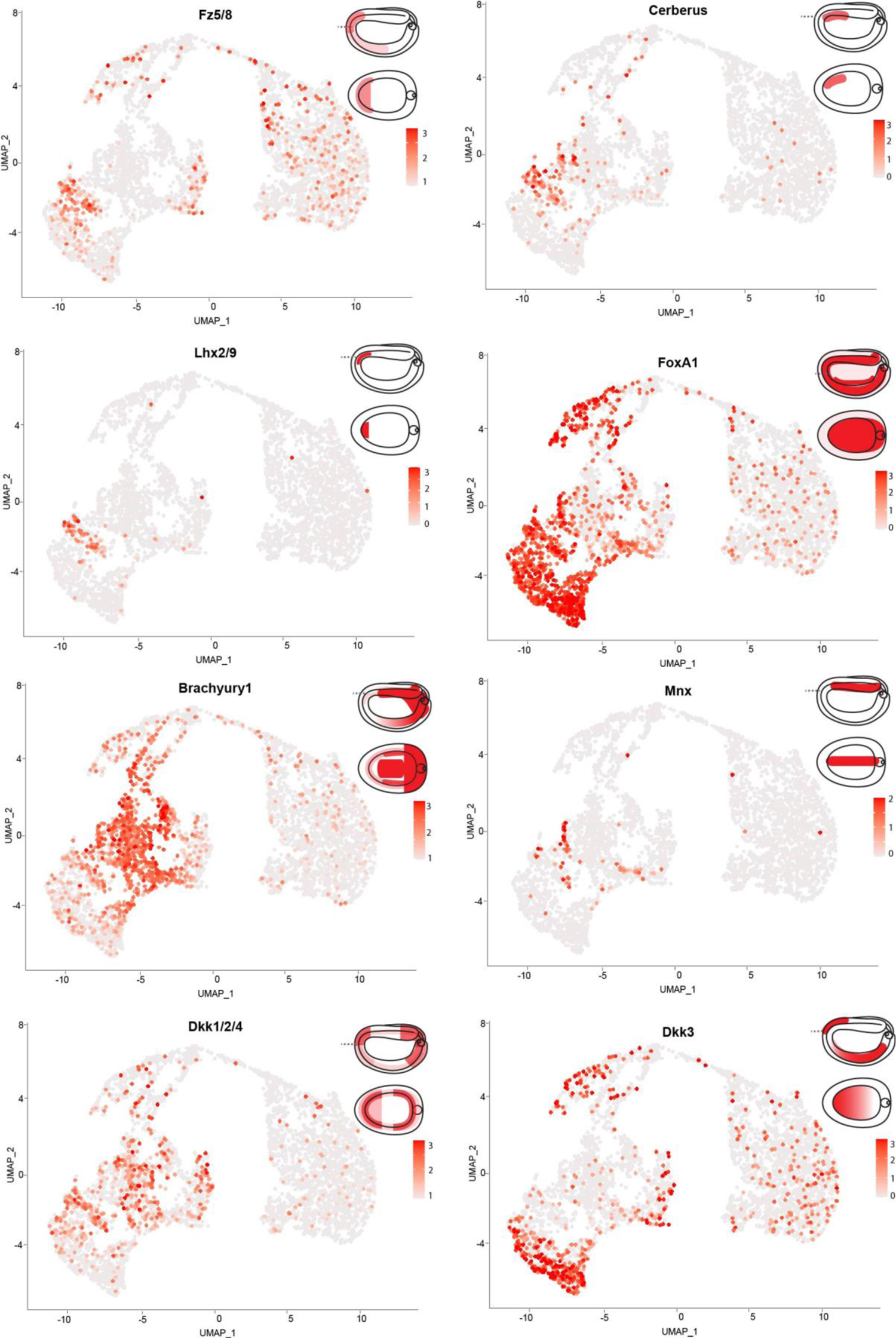
Feature plots of cells which are positive for each individual marker gene from Fig.**1*N0*** with schematic drawings of spatial gene expression (starting from the top: lateral view, dorsal view section trough the position marked with dashed line in lateral view).

**fig.S2d:**
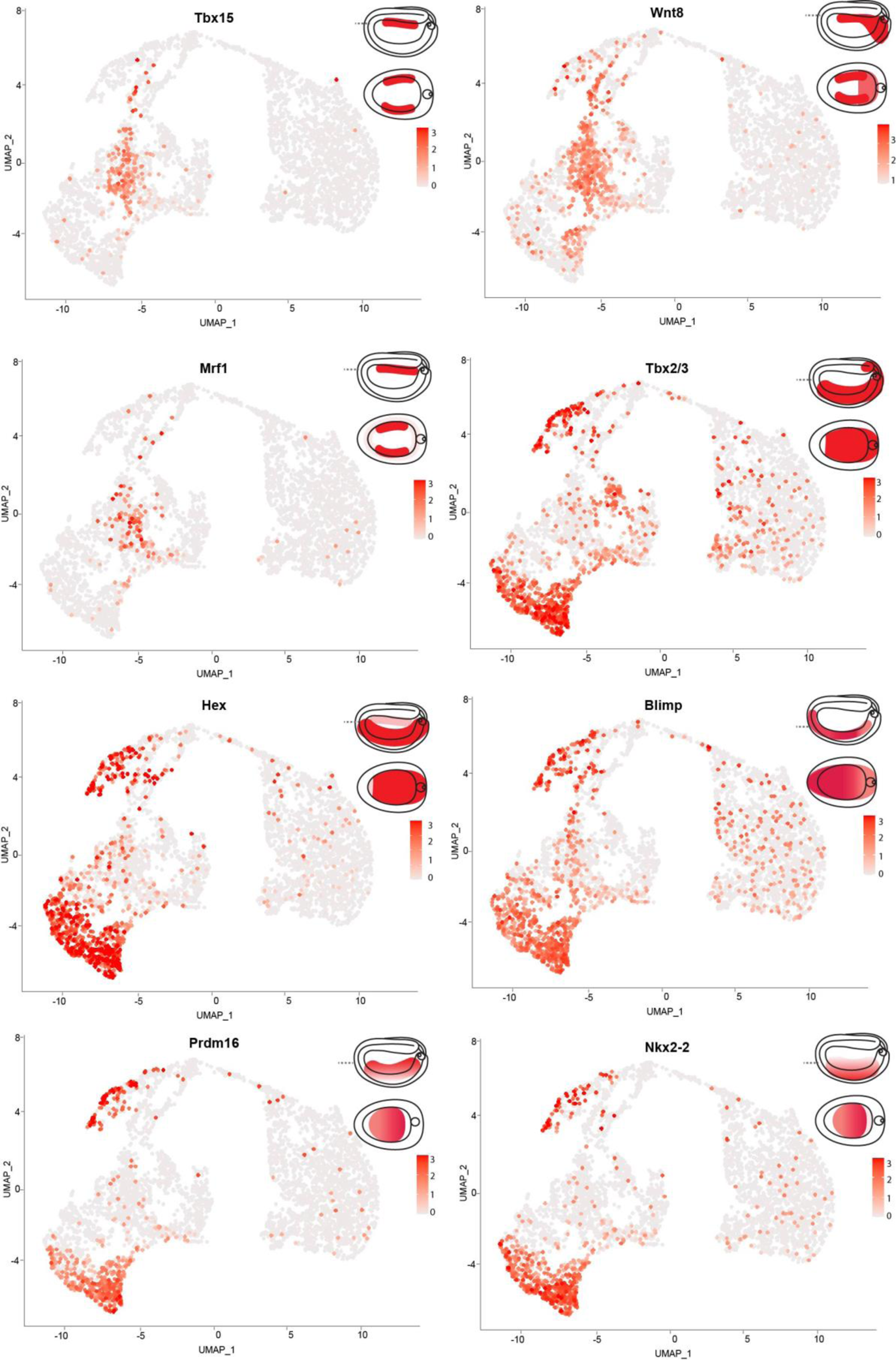
Feature plots of cells which are positive for each individual marker gene from Fig.**1*N0*** with schematic drawings of spatial gene expression (starting from the top: lateral view, dorsal view section trough the position marked with dashed line in lateral view).

**fig.S2e:**
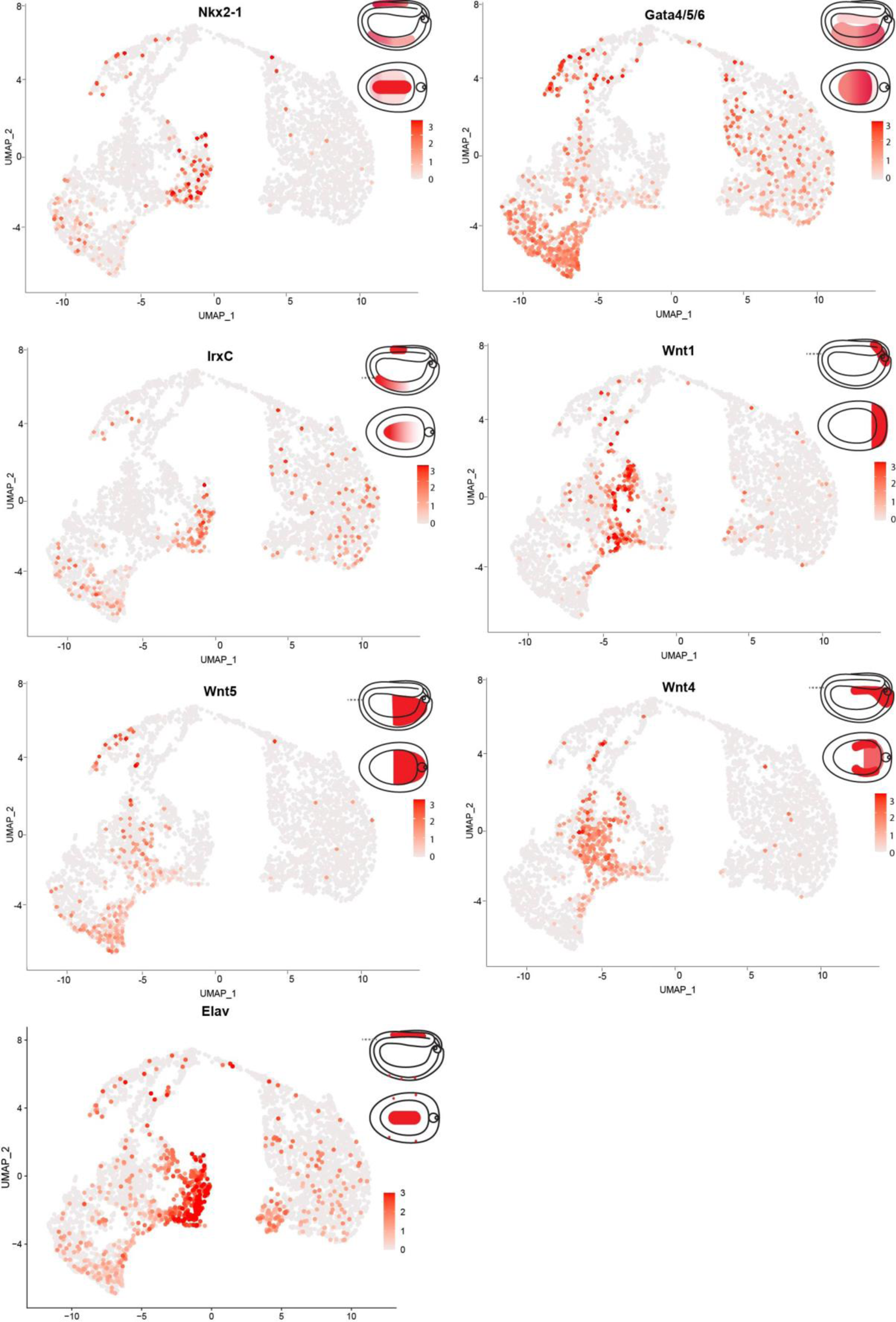
Feature plots of cells which are positive for each individual marker gene from Fig.**1*N0*** with schematic drawings of spatial gene expression (starting from the top: lateral view, dorsal view section trough the position marked with dashed line in lateral view).

**fig.S3:**
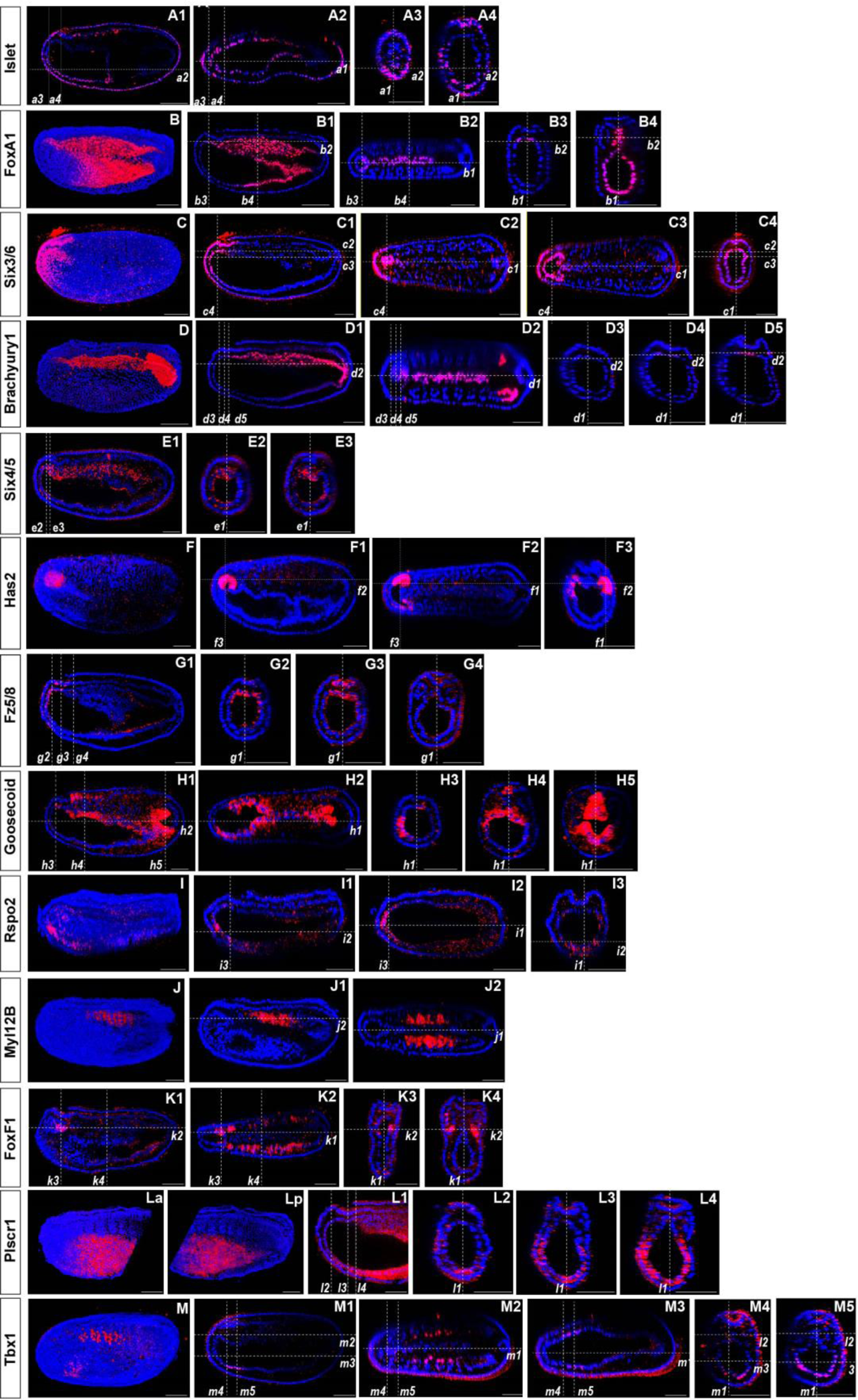
Spatial gene expression of individual genes at the N2 stage. The position of individual z-slices (**A1-A4, B1-B4, C1-C4, D1-D5, E1-E3, F1-F3, G1-G4, H1-H5, I1-I3, J1-J2, K1-K4, L1-L4, M1-M5**) from complete z-stacks are indicated with dashed lines (*a1-i4, b1-b4, c1-c4, d1-d5, e1-e3, f1-f3, g1-g4, h1-h5, i1-i3, j1-j2, k1-k4, l1-l4, m1-m5*). (**A, B, C, D, F, I, J, La** (anterior part)**, Lp** (posterior part**), M**) Lateral 3D views of substacks from a complete z-stacks.

**fig.S3a:**
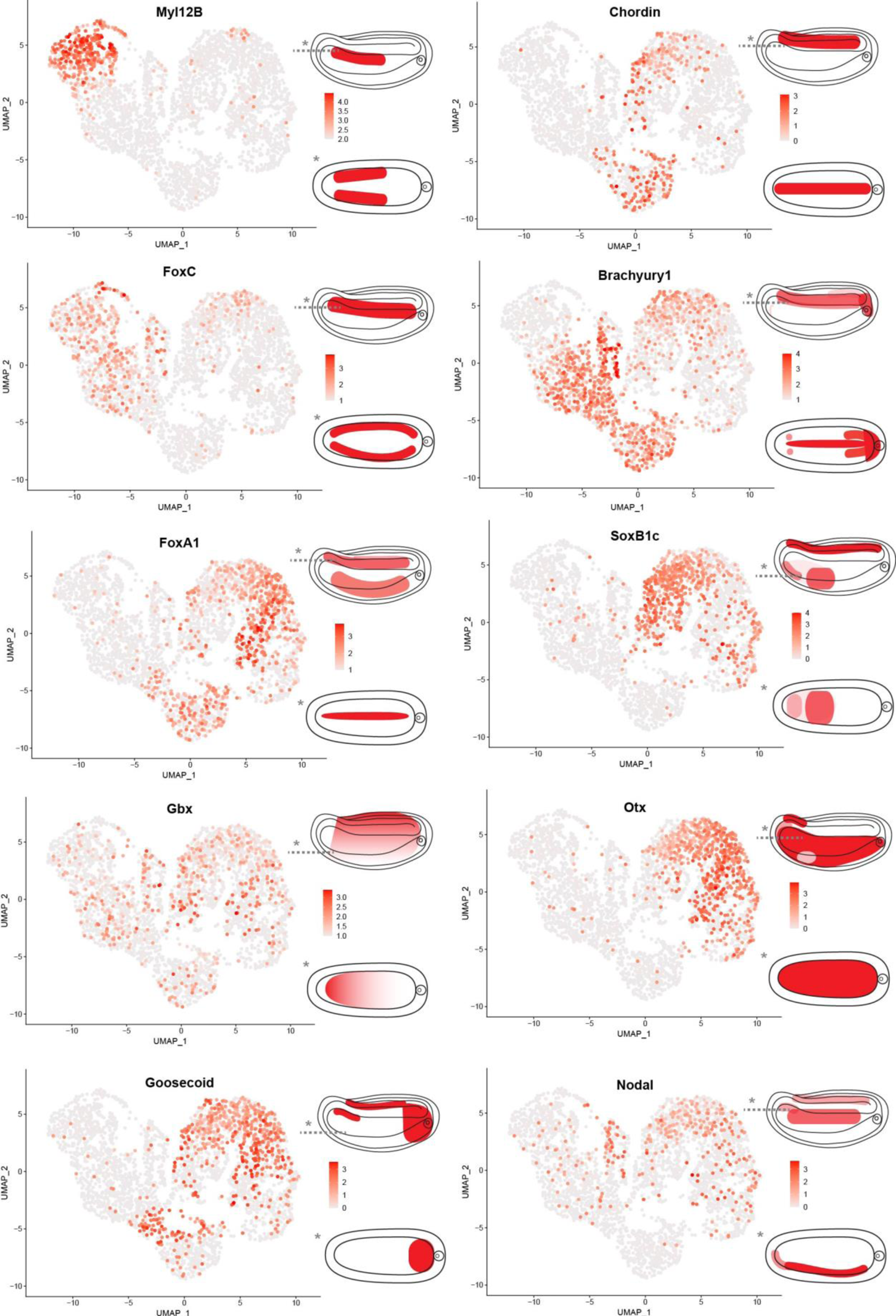
Feature plots of cells which are positive for each individual marker gene from Fig.**1*N2*** with schematic drawings of spatial gene expression (starting from the top: lateral view, dorsal view section

**fig.S3b:**
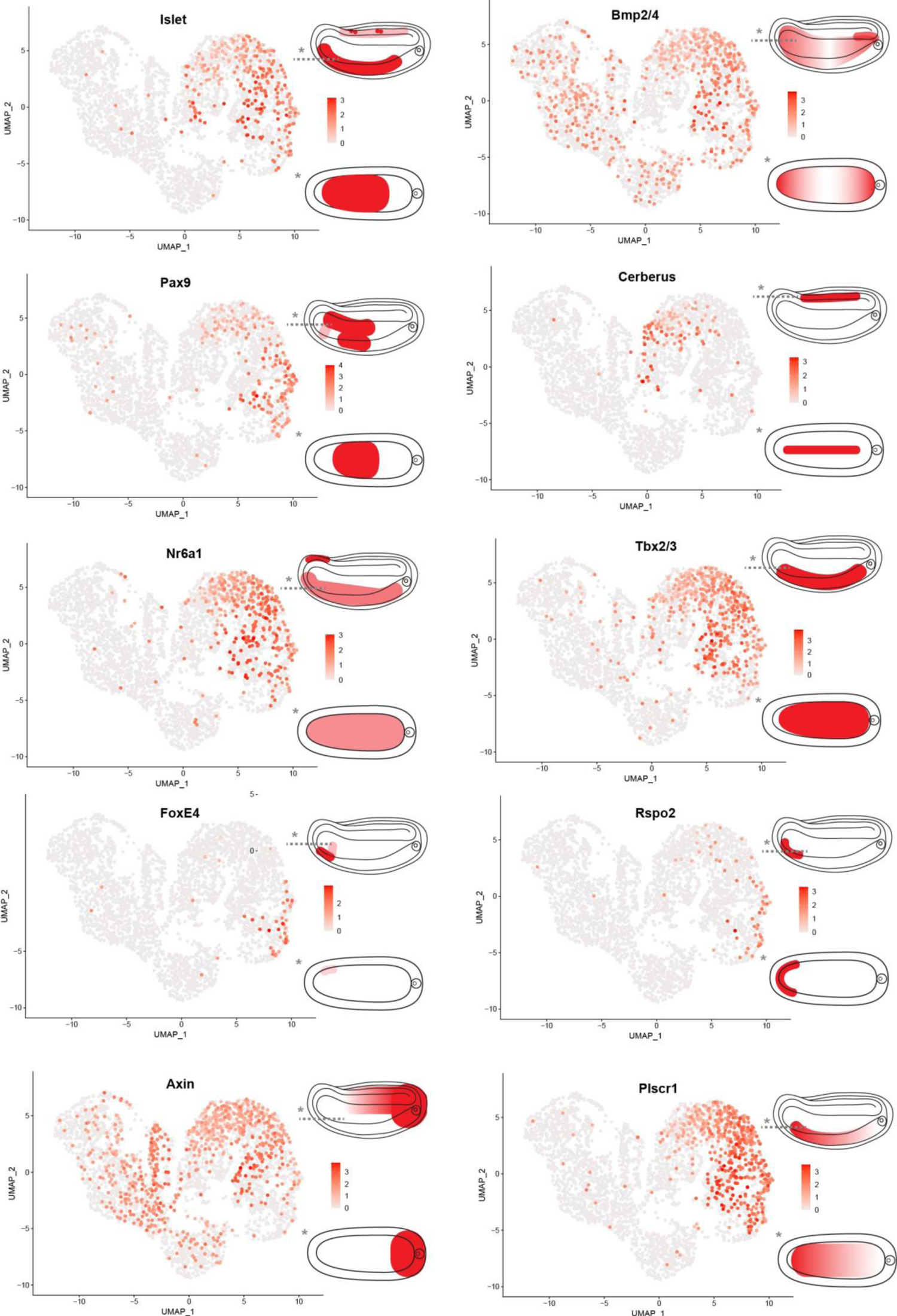
Feature plots of cells which are positive for each individual marker gene from Fig.**1*N2*** with schematic drawings of spatial gene expression (starting from the top: lateral view, dorsal view section

**fig.S3c:**
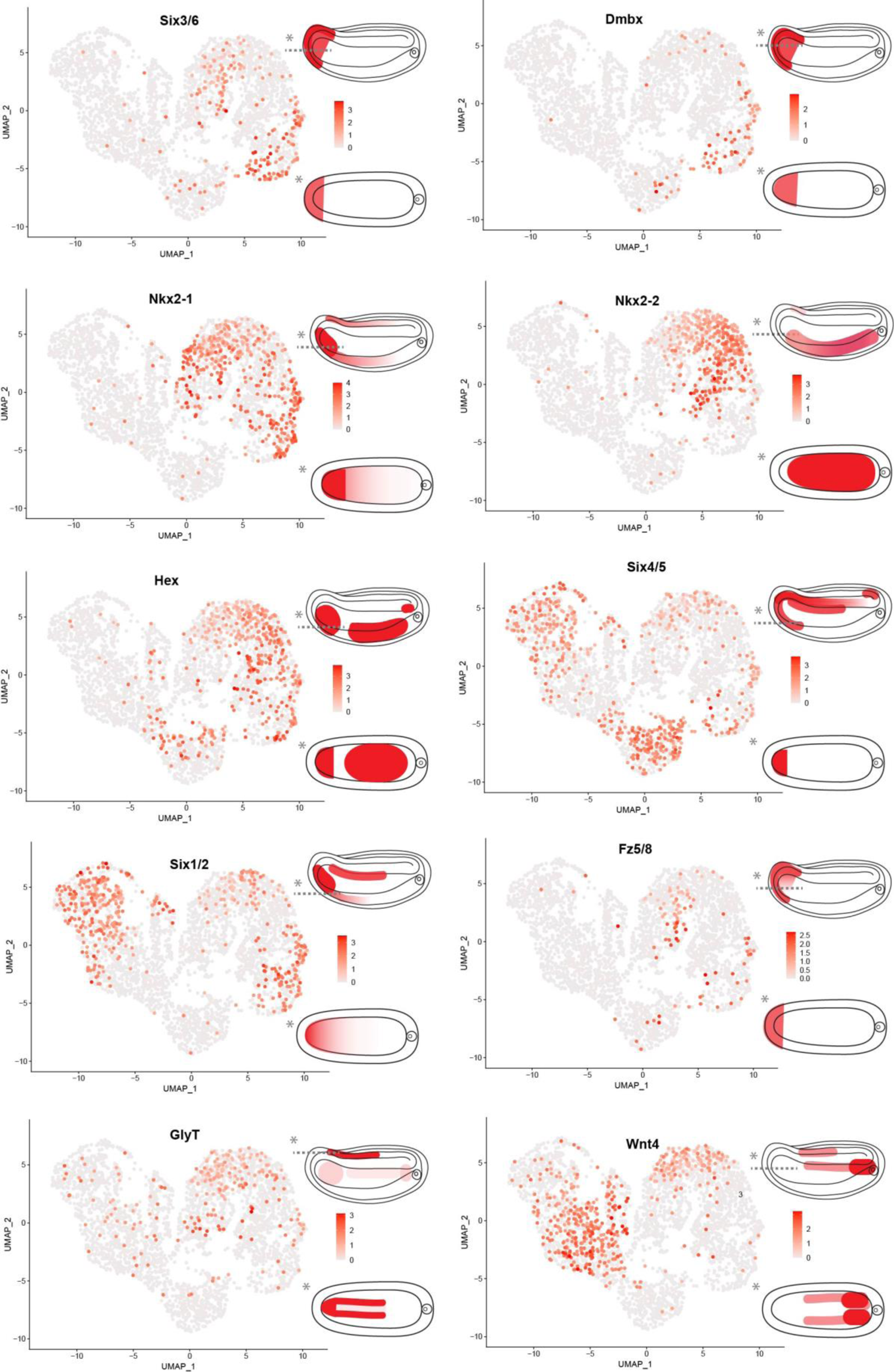
Feature plots of cells which are positive for each individual marker gene from Fig.**1*N2*** with schematic drawings of spatial gene expression (starting from the top: lateral view, dorsal view section trough the position marked with dashed line in lateral view).

**fig.S3d:**
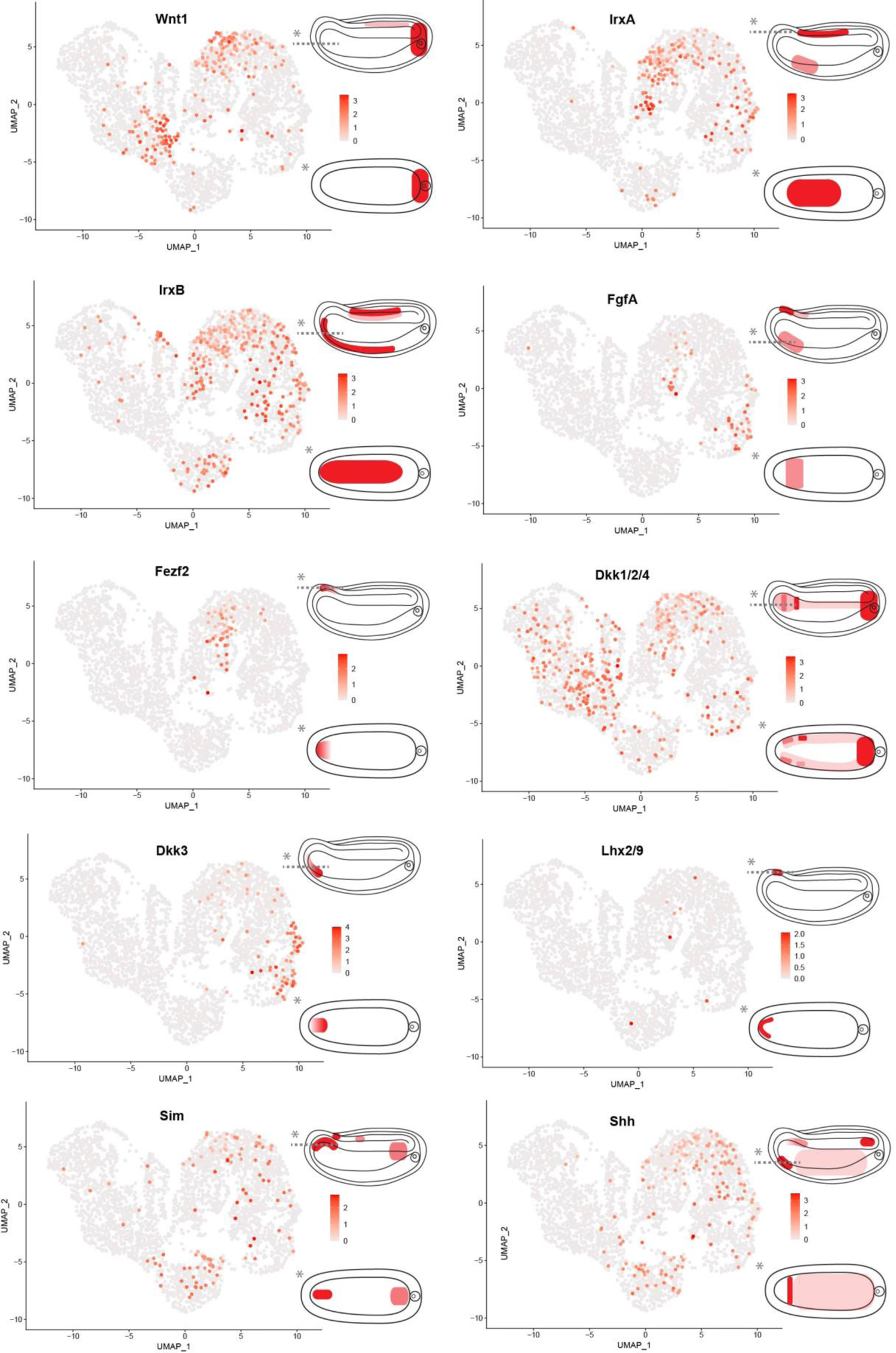
Feature plots of cells which are positive for each individual marker gene from Fig.**1*N2*** with schematic drawings of spatial gene expression (starting from the top: lateral view, dorsal view section trough the position marked with dashed line in lateral view).

**fig.S3e:**
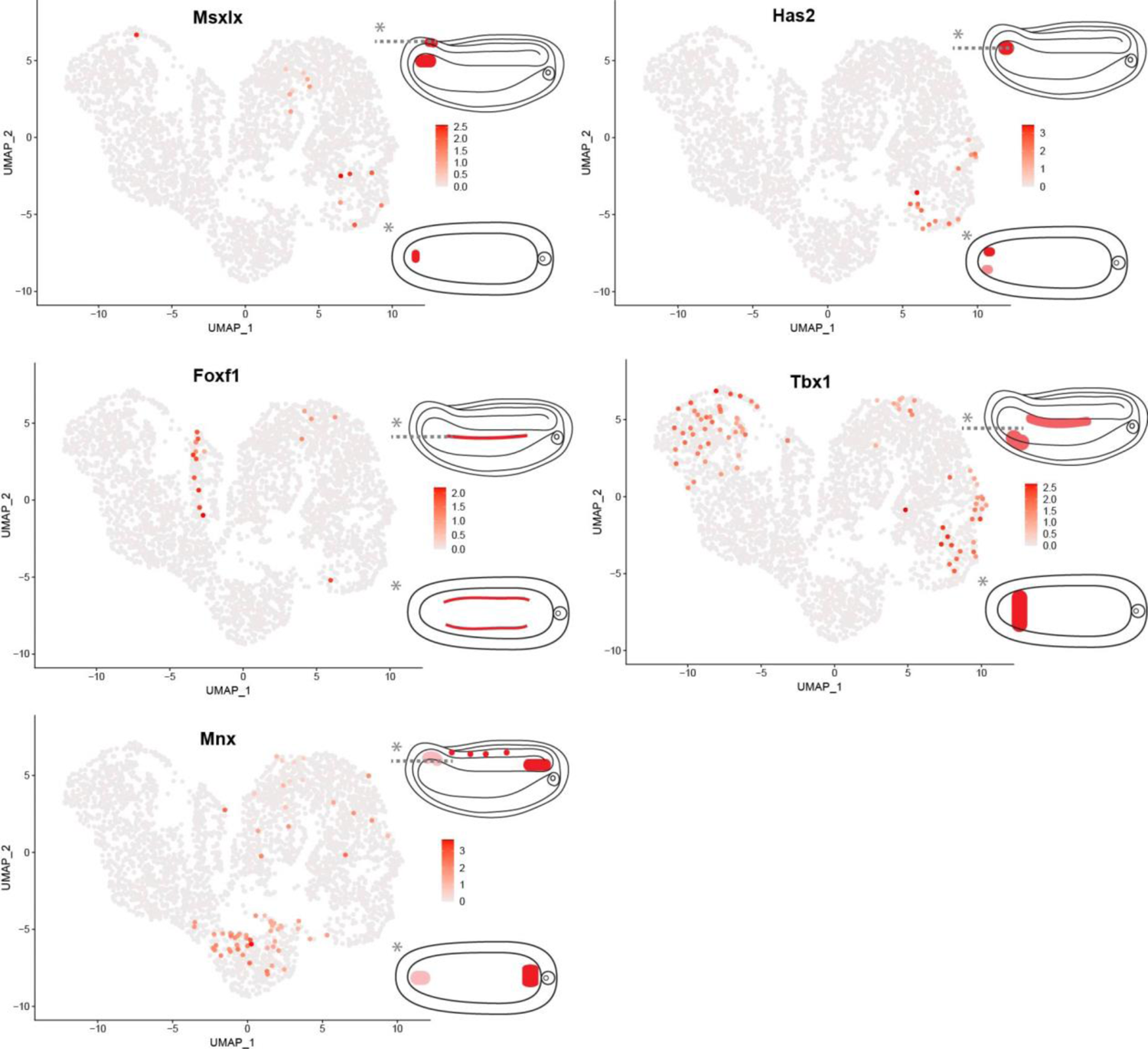
Feature plots of cells which are positive for each individual marker gene from Fig.**1*N2*** with schematic drawings of spatial gene expression (starting from the top: lateral view, dorsal view section trough the position marked with dashed line in lateral view).

**fig.S4:**
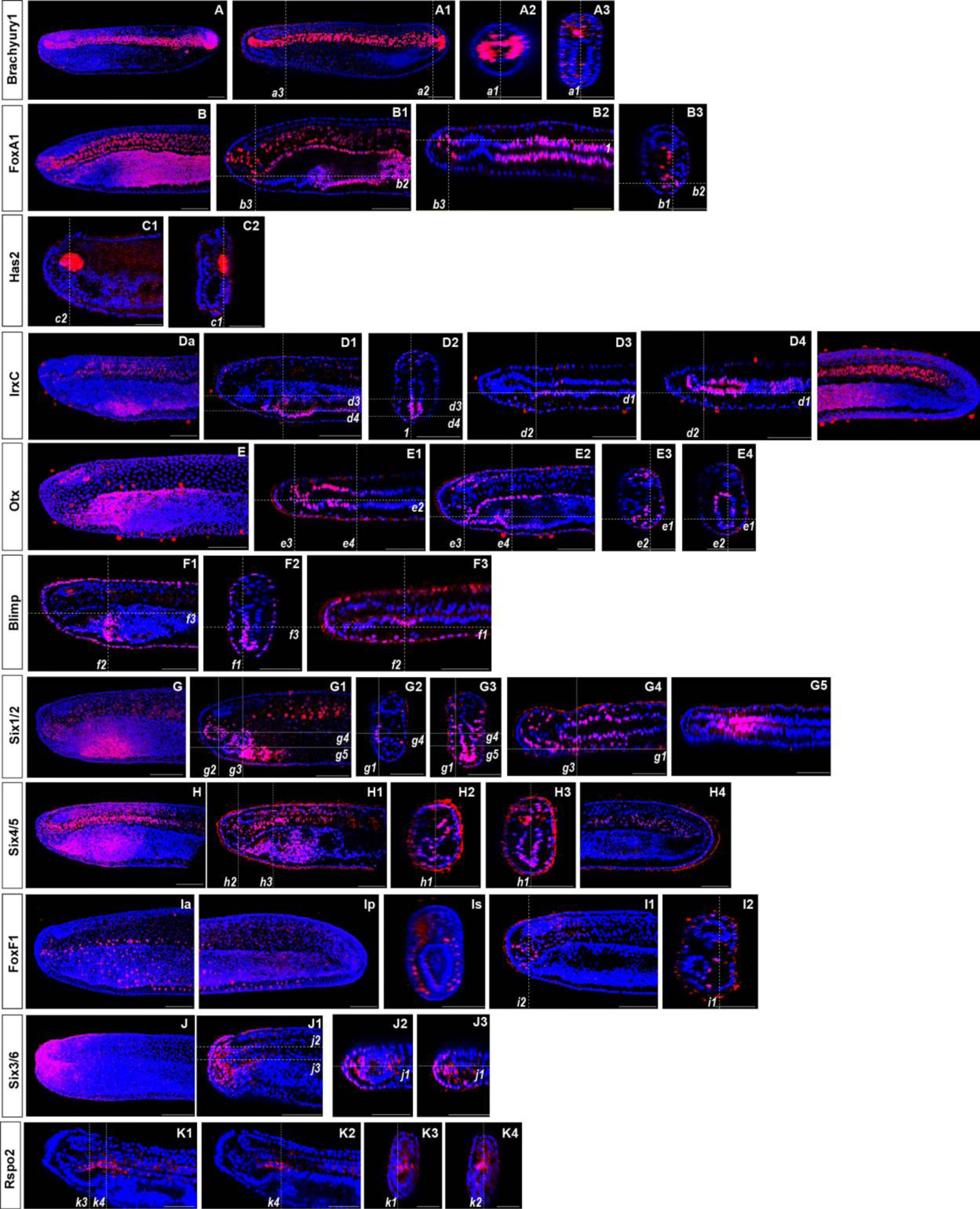
Spatial gene expression of individual genes at the N5 stage. The position of individual z-slices (**A1-A4, B1-B3, C1-C4, D1-D4, E1-E4, F1-F3, G1-G5, H1-H4, I1-I2, J1-J2, K1-K4, L1-L2**) from complete z-stacks are indicated with dashed lines (*a1-a43, b1-b3, c1-c4, d1-d4, e1-e4, f1-f3, g1-g5, h1-h4, i1-i2, j1-j2, k1-k4, l1-l2*). Lateral 3D views of substacks from a complete z-stacks in (**A, B, Da, J** (anterior part)**, Dp** (posterior part)**, E, G, H, Ia, Ip, Is** (transverse section view).

**fig.S4a:**
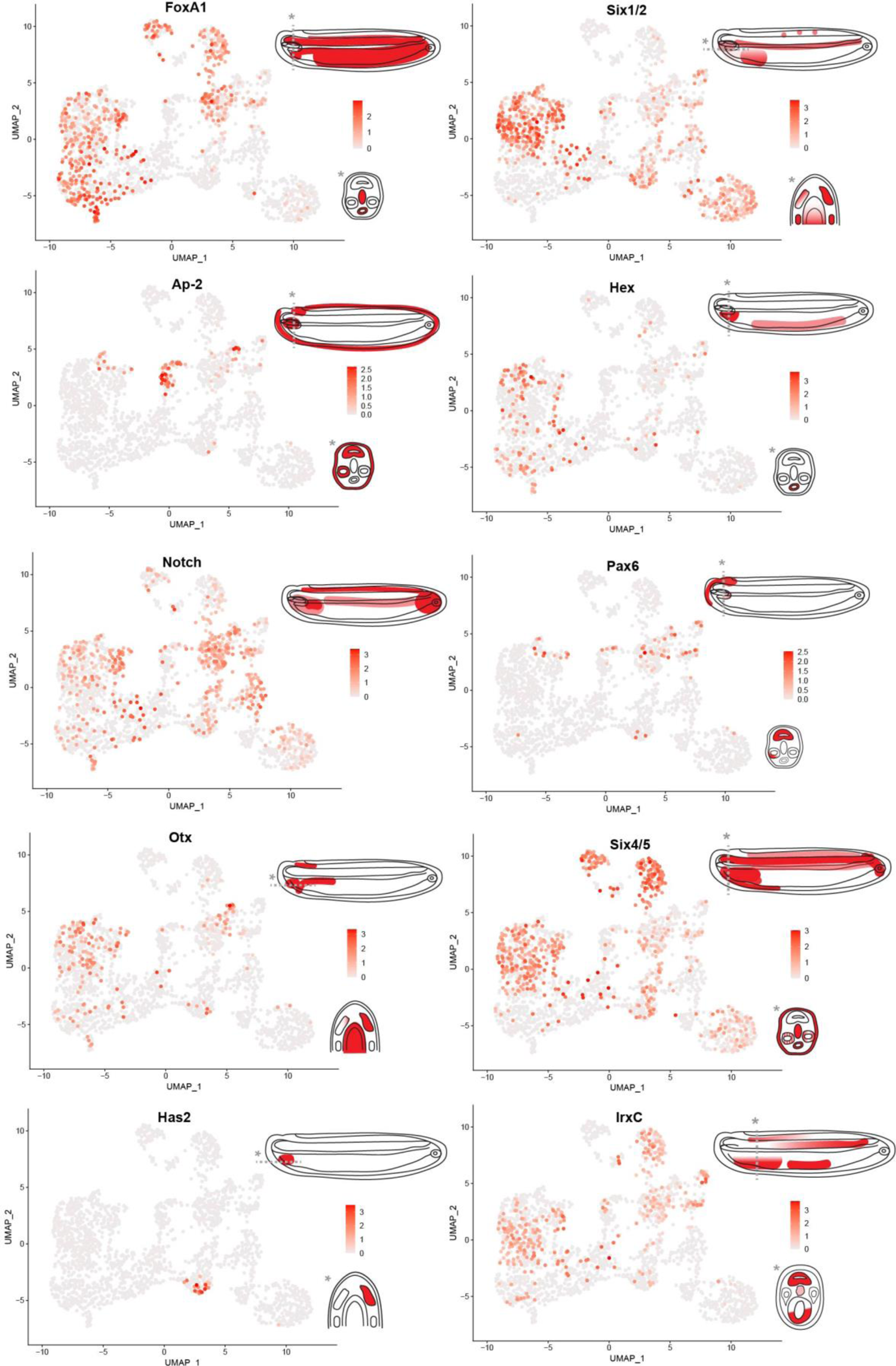
The feature plots showcase cells that are positive for each individual marker gene from Fig.**1*N5***, accompanied by schematic drawings of spatial gene expression (starting from the top: lateral view, the

**fig.S4b:**
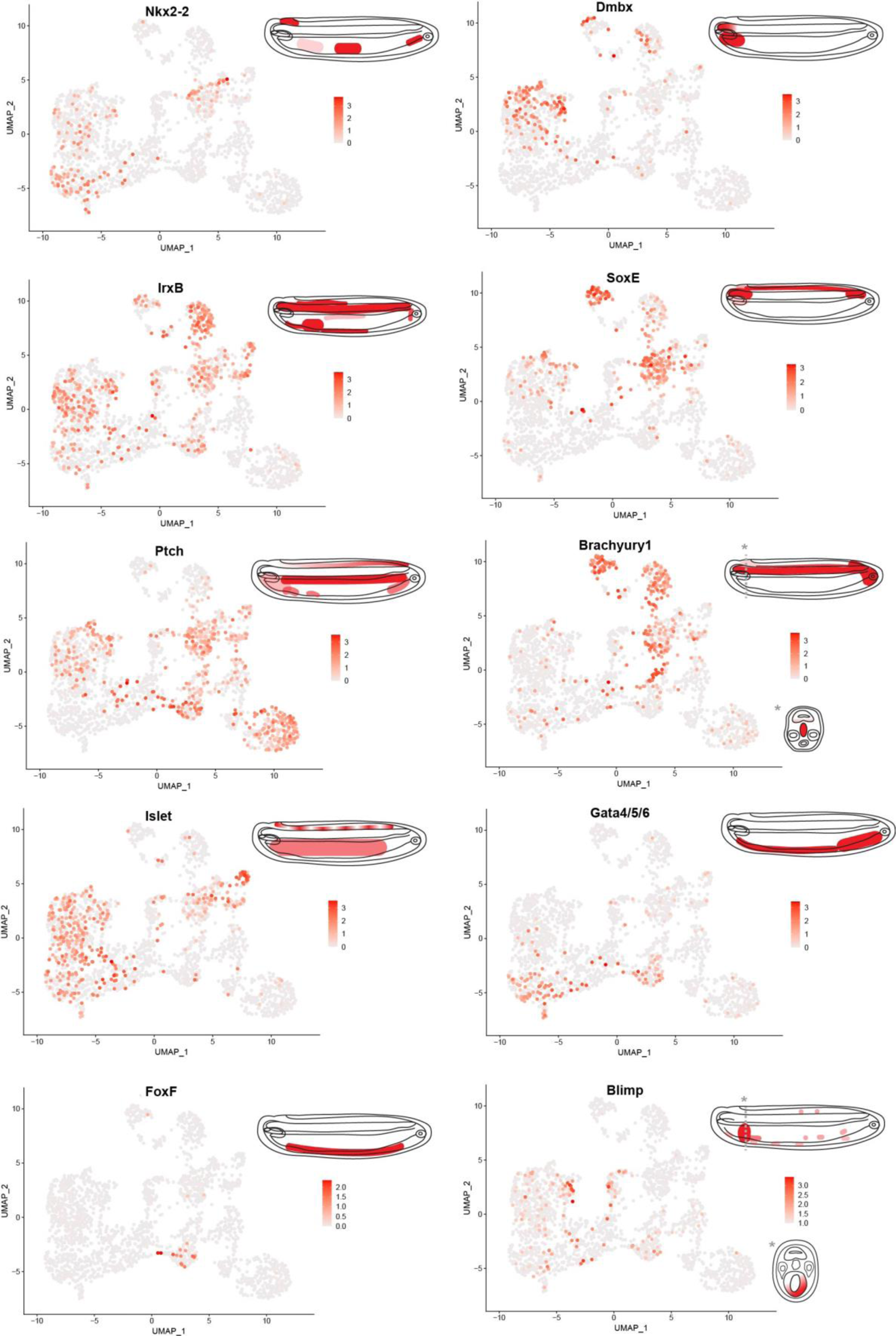
The feature plots showcase cells that are positive for each individual marker gene from Fig.**1*N5***, accompanied by schematic drawings of spatial gene expression (starting from the top: lateral view, the

**fig.S4c:**
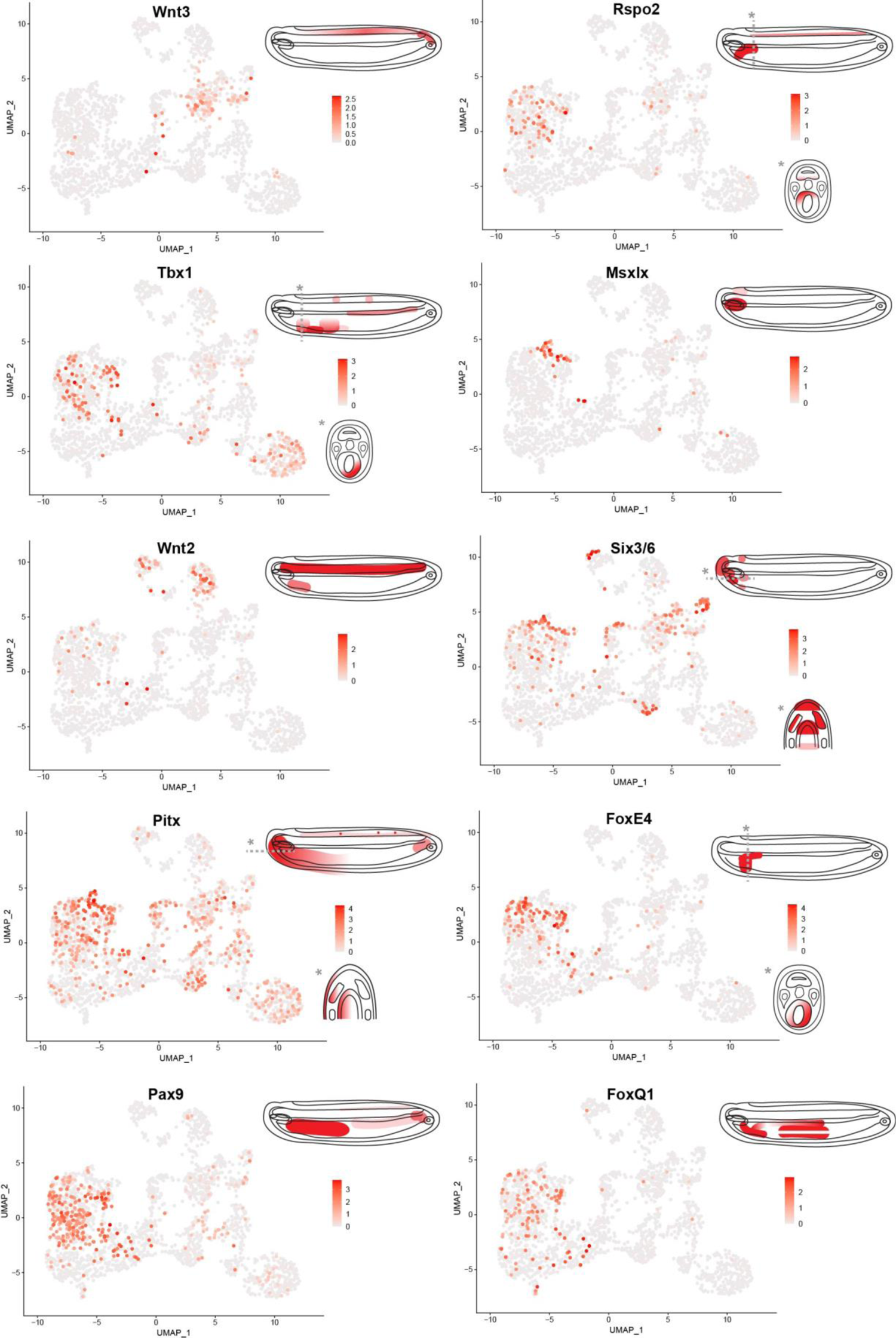
The feature plots showcase cells that are positive for eachindividual marker gene from Fig.**1*N5***, accompanied by schematic drawings of spatial gene expression (starting from the top: lateral view, the section trough the position marked with dashed line in lateral view).

**fig.S4d:**
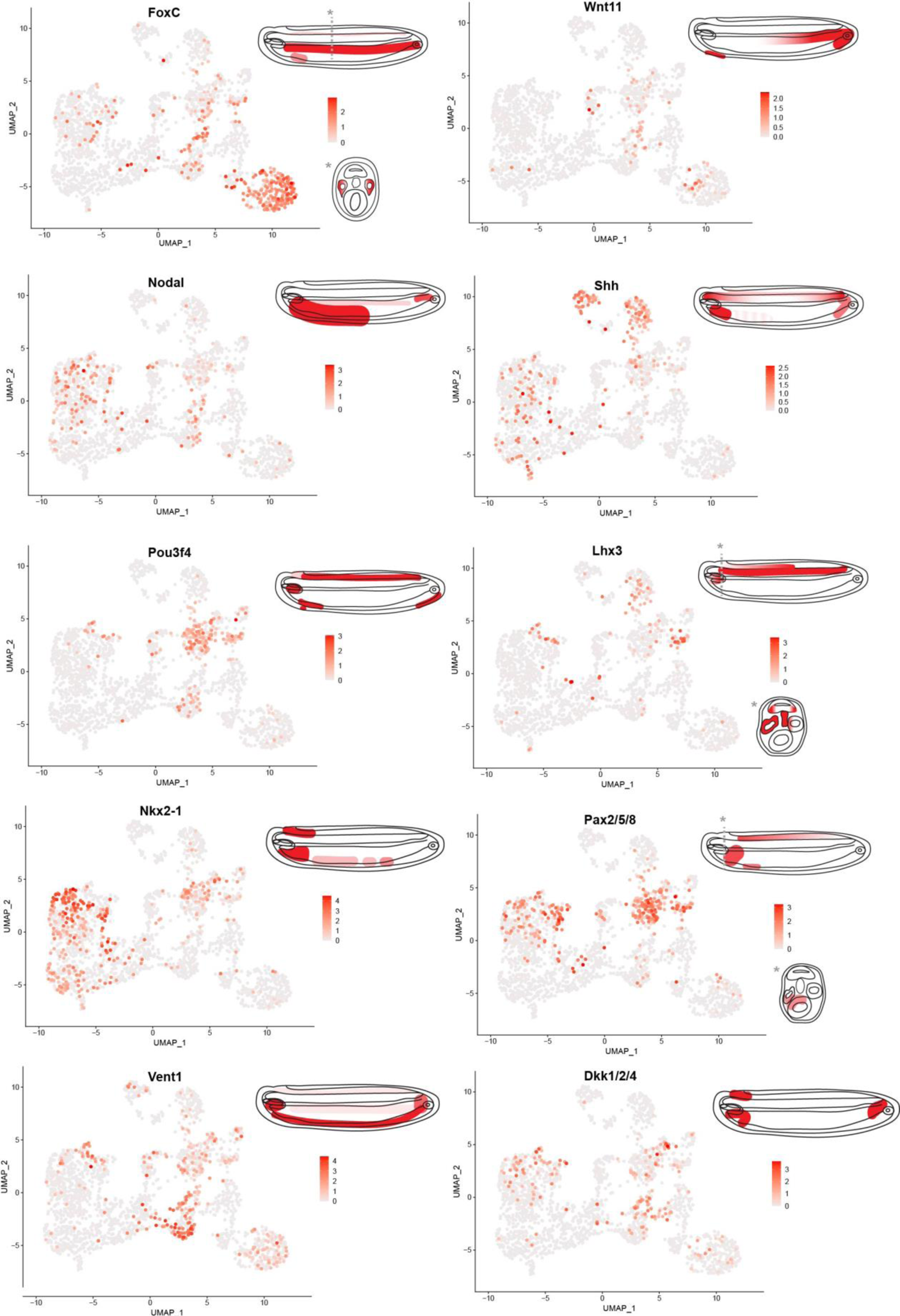
The feature plots showcase cells that are positive for each individual marker gene from Fig.**1*N5***, accompanied by schematic drawings of spatial gene expression (starting from the top: lateral view, the section trough the position marked with dashed line in lateral view).

**fig.S4e:**
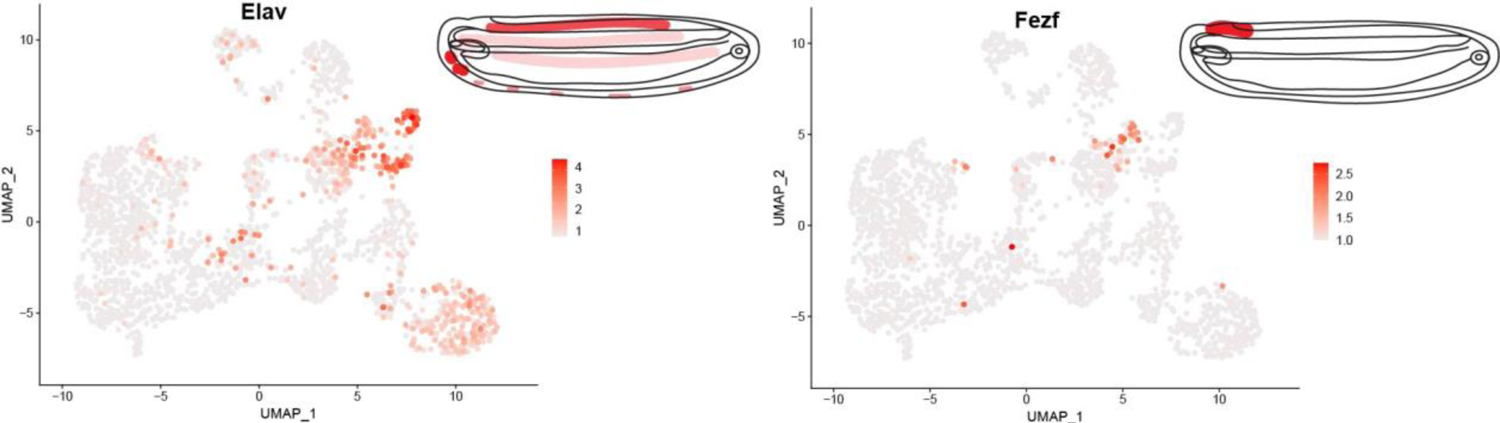
The feature plots showcase cells that are positive for each individual marker gene from Fig.**1*N5***, accompanied by schematic drawings of spatial gene expression.

**fig.S5:**
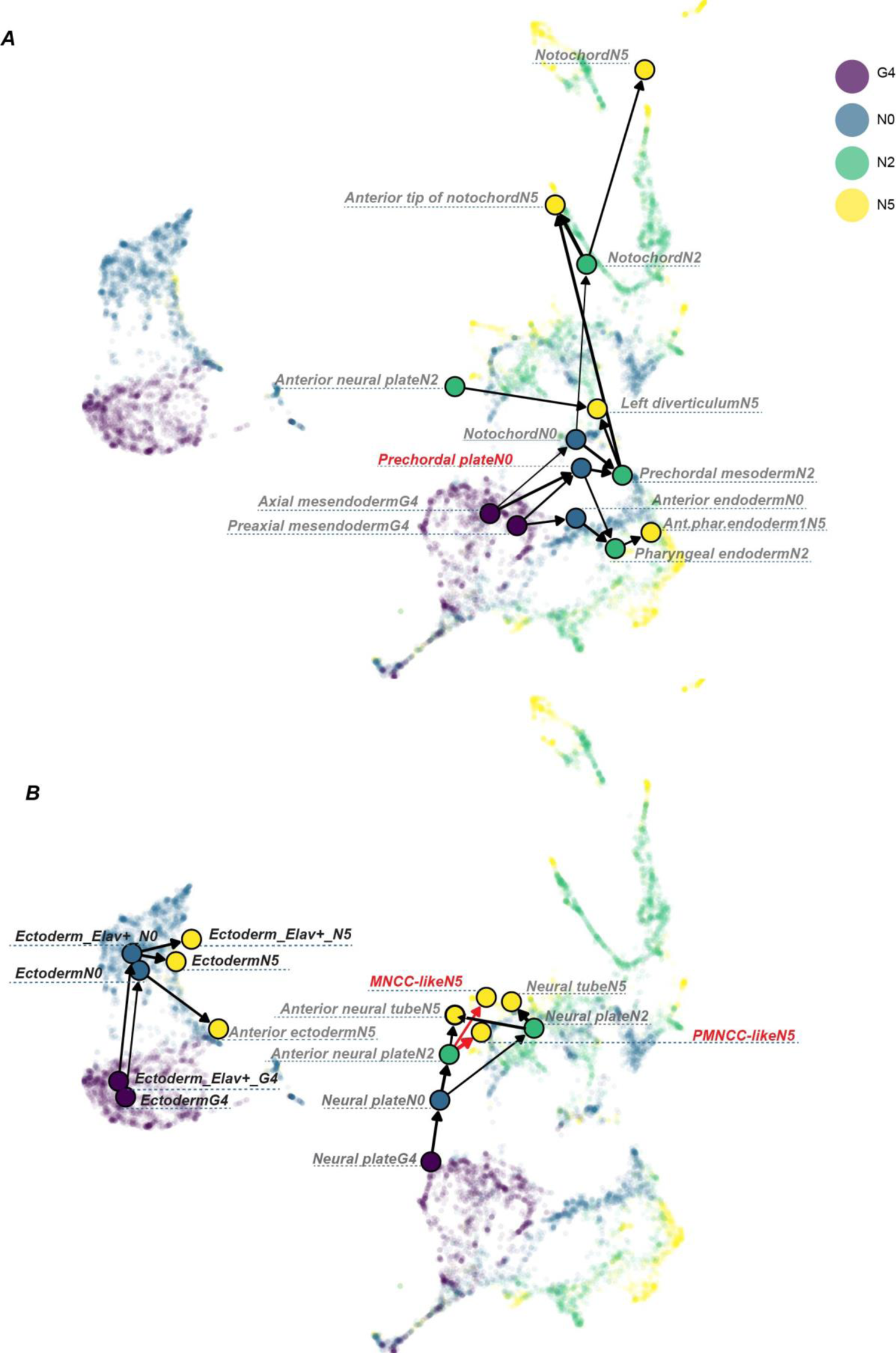
Cell fate transition probabilities across the G4, N0, N2 and N5 stages. Scaled transition probabilities of select cell populations, centering on prechordal plate (***A***) and MNCC-like (***B***), are visualized through denoised directed graphs superimposed on a UMAP of the MNN-integrated dataset. The width of the edges corresponds to the edge weight, representing the scaled probability.

**fig.S6:**
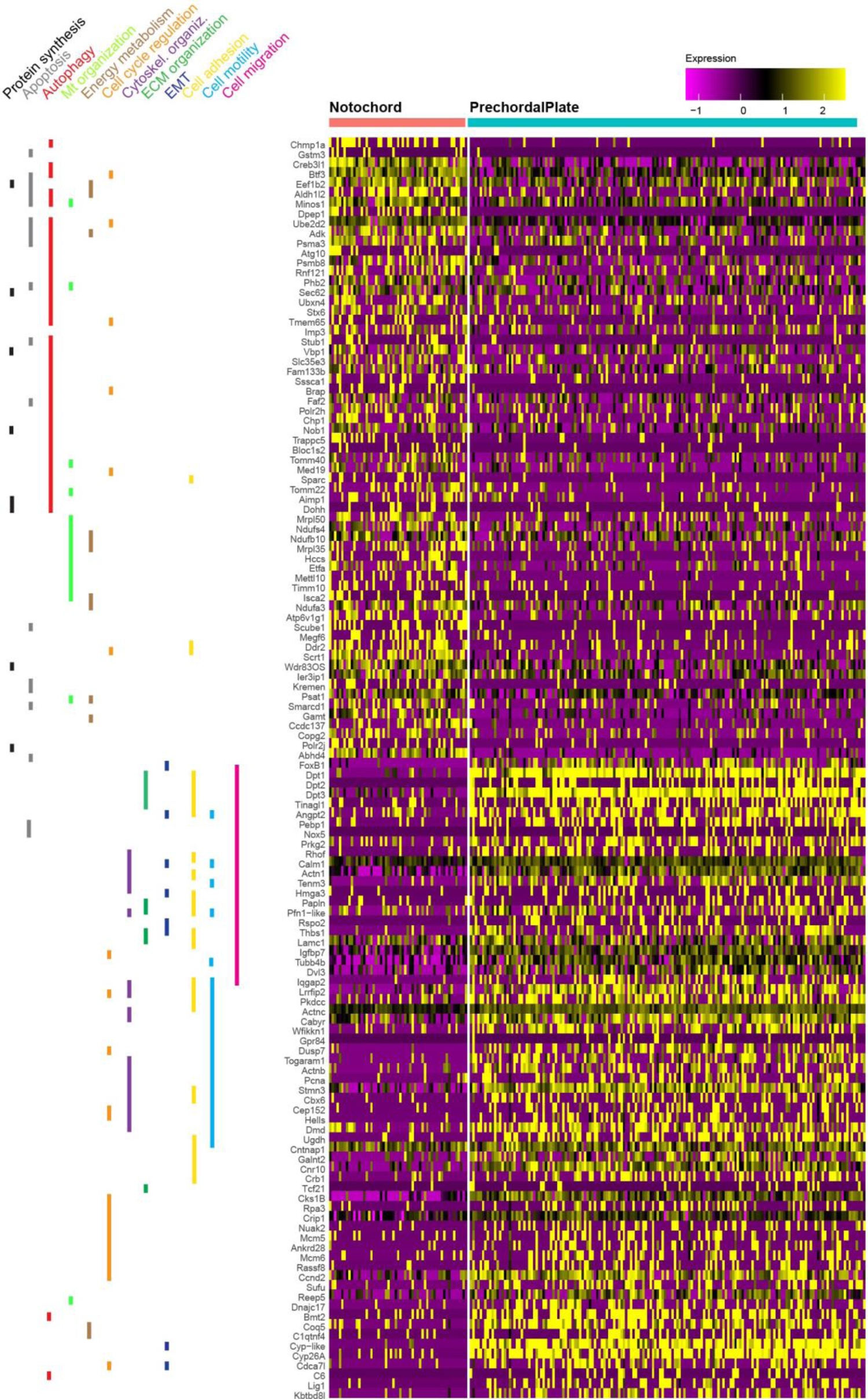
Heat map of single-cell expression data showing the differentially expressed genes specific to the prechordal plate and notochord clusters at the N0 stage. Cells are displayed in columns, and genes are represented in rows. Colored vertical bars located to the right of the gene names indicate the cellular processes in which the genes might be involved.

**fig.S7:**
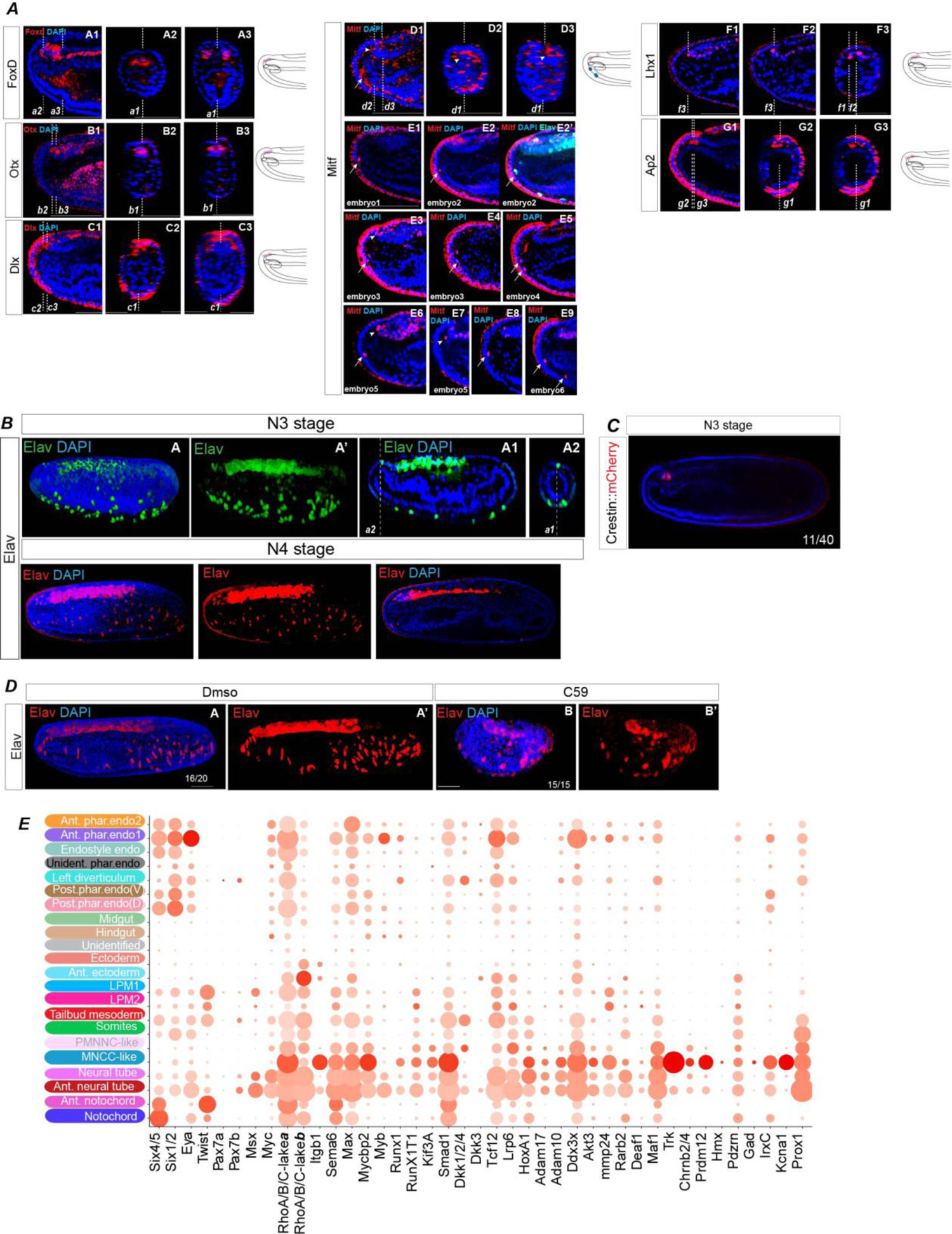
Neural crest-like cells are present in amphioxus. (***A***) The expression of FoxD, Otx, Dlx, Lhx1 and Ap-2 in PMNCC, and Mitf in PMNCC and MNCC in the context of the embryo at N5. Arrows in (***A*D1-E9**) indicate the expression in MNCC, while arrowheads in the cells of the rostral cerebral vesicle. mRNA is detected in (**A1-D3**, **F1-G3**) and protein in (**E1-E9**) (***B***) Elav expression at N3 and N4. The position of individual stacks (***A*A1-D3**, ***A*F1-G3, *B*A1-A2**) are indicated with dashed lines and marked as ***a1-d3***, ***f1-g3***, ***a1-a2***. (***C***) The activity of Crestin::mCherry in the embryos at N3. (***D***) Wnt signaling inhibition blocks the emergence of Elav+ MNCC. (***E***) The DotPlot illustrates DEG within PMNCC-like and MNCC-like populations.

**fig.S8:**
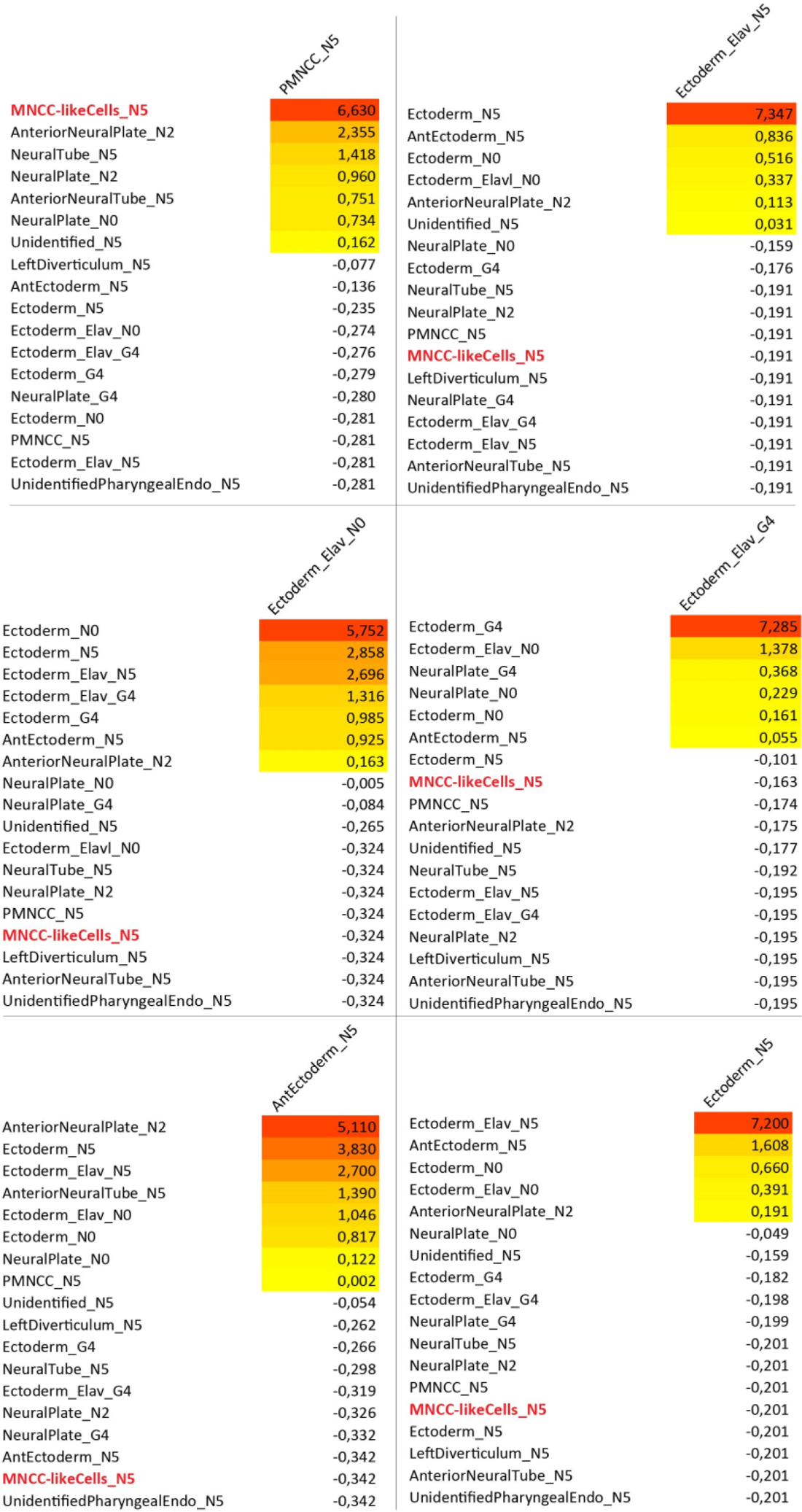
The probabilities of MNCC-like cell originating from PMNCC or ectoderm. Transition probabilities are determined by the Euclidean distance between cells in gene expression space, calculated as a diffusion map. The transition probabilities are biased by the differences in each cell’s assigned pseudotime, favoring transitions that align with the direction of real time. Values represent z-scaled transition probabilities of cell type in column’s header transiting to cell types in rows. Cell types are divided by timepoints.

**fig.S9:**
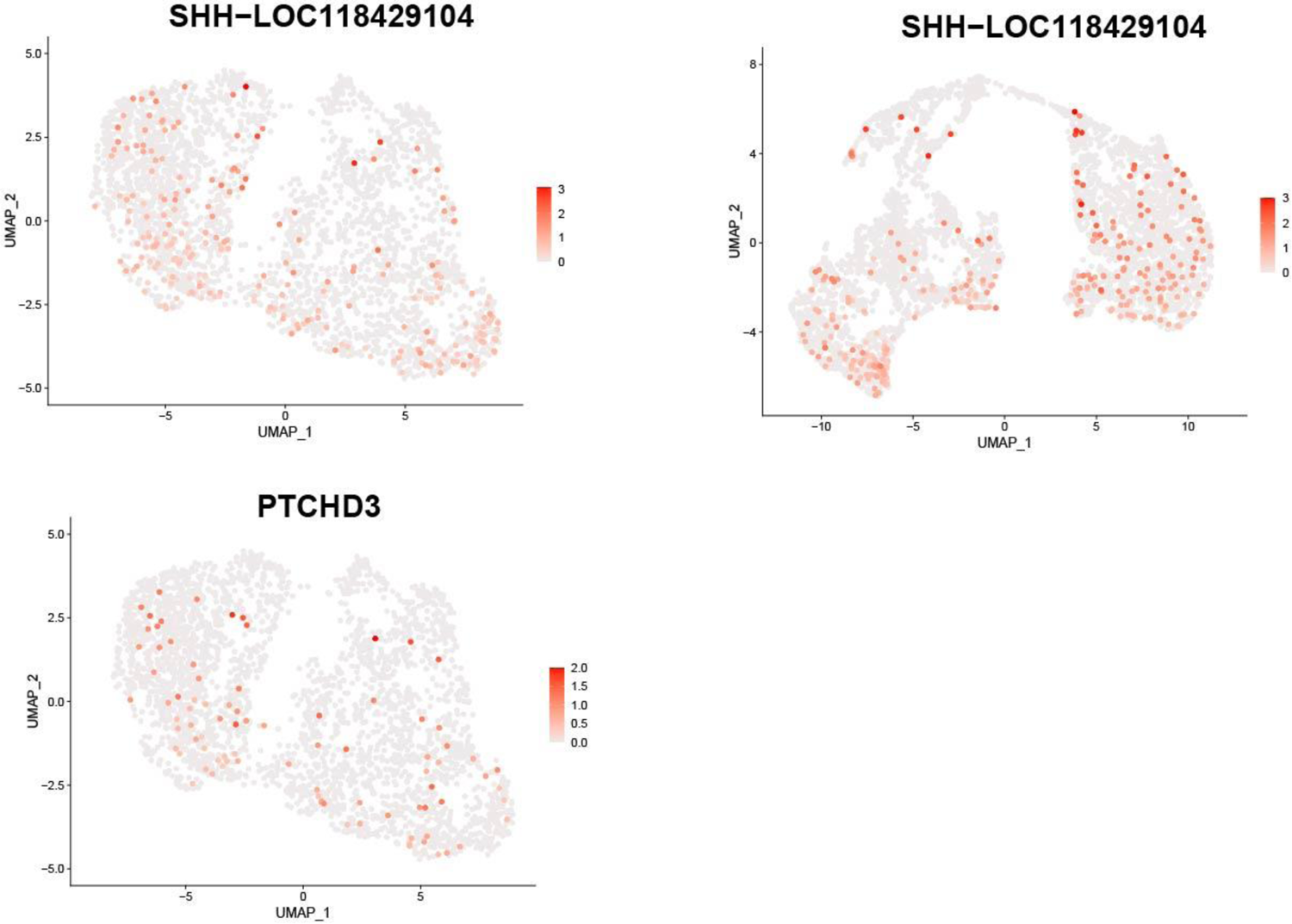
Patched/Shh-like genes are expressed at the G4 and N0 stages.

**fig.S10:**
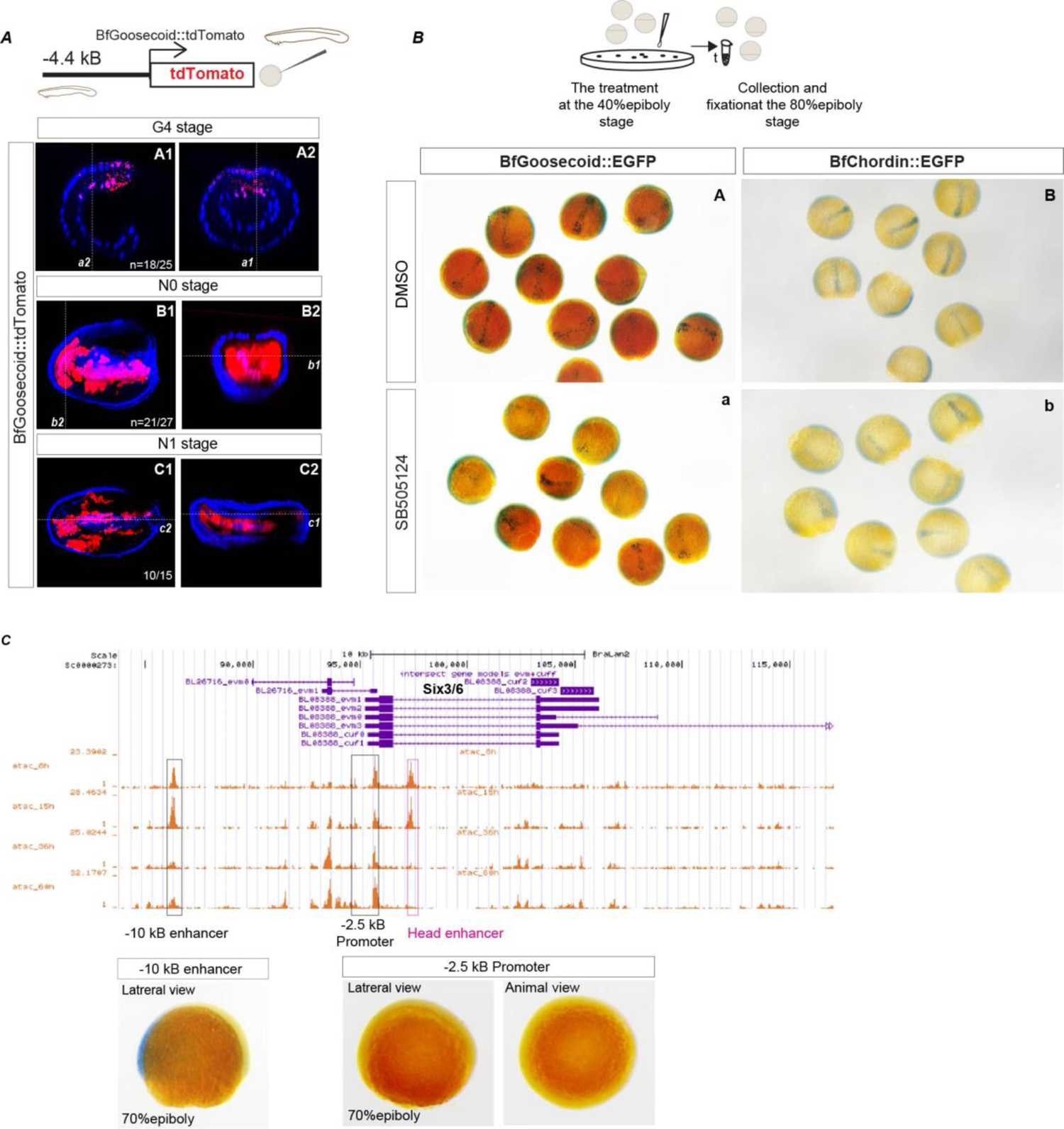
Homology in chordate prechordal plate development at the cis-regulatory level. (***A***) The activity of amphioxus Goosecoid cis-regulatory sequence in transient transgenic amphioxus embryos. (**A*B1-C2***) 3D-views of substacks from z-stacks. Z-slices of the whole embryo in (**A*A1-A2***). Dorsal (**A*B1***, **A*C1***), lateral (**A*C2***) and frontal (**A*B2***) views. Dashed lines in (***A***) indicate positions of sections. (***B***) Inhibition of Nodal signaling with SB505124 at the early gastrula stage downregulates the activity of BfGoosecoid::EGFP (***B*A-a**), Bf::Chordin-EGFP (***B*B-b**) in the prechordal plate region of zebrafish. (***C***) Genome browser excerpt showing open chromatin regions within Six3/6 locus as determined by ATAC-seq. The putative regulatory regions tested functionally by transgenic experiments in the zebrafish are shown in brackets.

**fig.S11:**
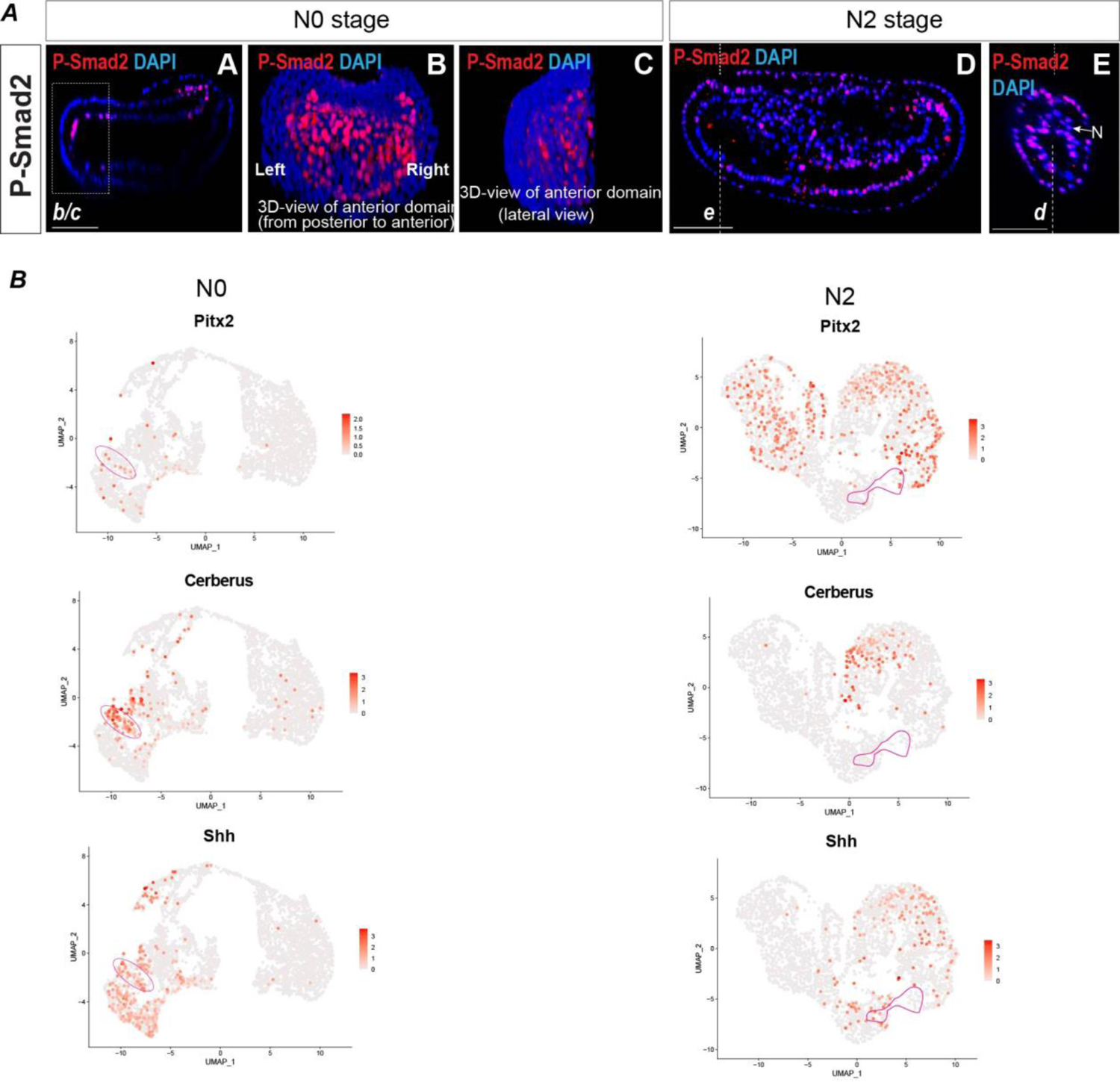
The activity of Nodal signaling and the expression of PrCP-specific genes during early neurula stages in amphioxus. (***A***) P-Smad2 immunostaining at N0 and at N2. (**A, D, E**) Individual stacks. Dashed rectangle (***b/c***) in (**A**) depicts the area shown in (**B**) and (**C**) as 3D-reconstruction. Dashed lines in (**D**) and (**E**) indicate the positions of individual stacks marked as (***e, d***). N – notochord. (***B***) Feature plots of Pitx2, Cerberus and Shh expression at N0 and N2. Pink closed lines indicate PrCP at N0 and prechordal mesoderm at N2.

**fig.S12:**
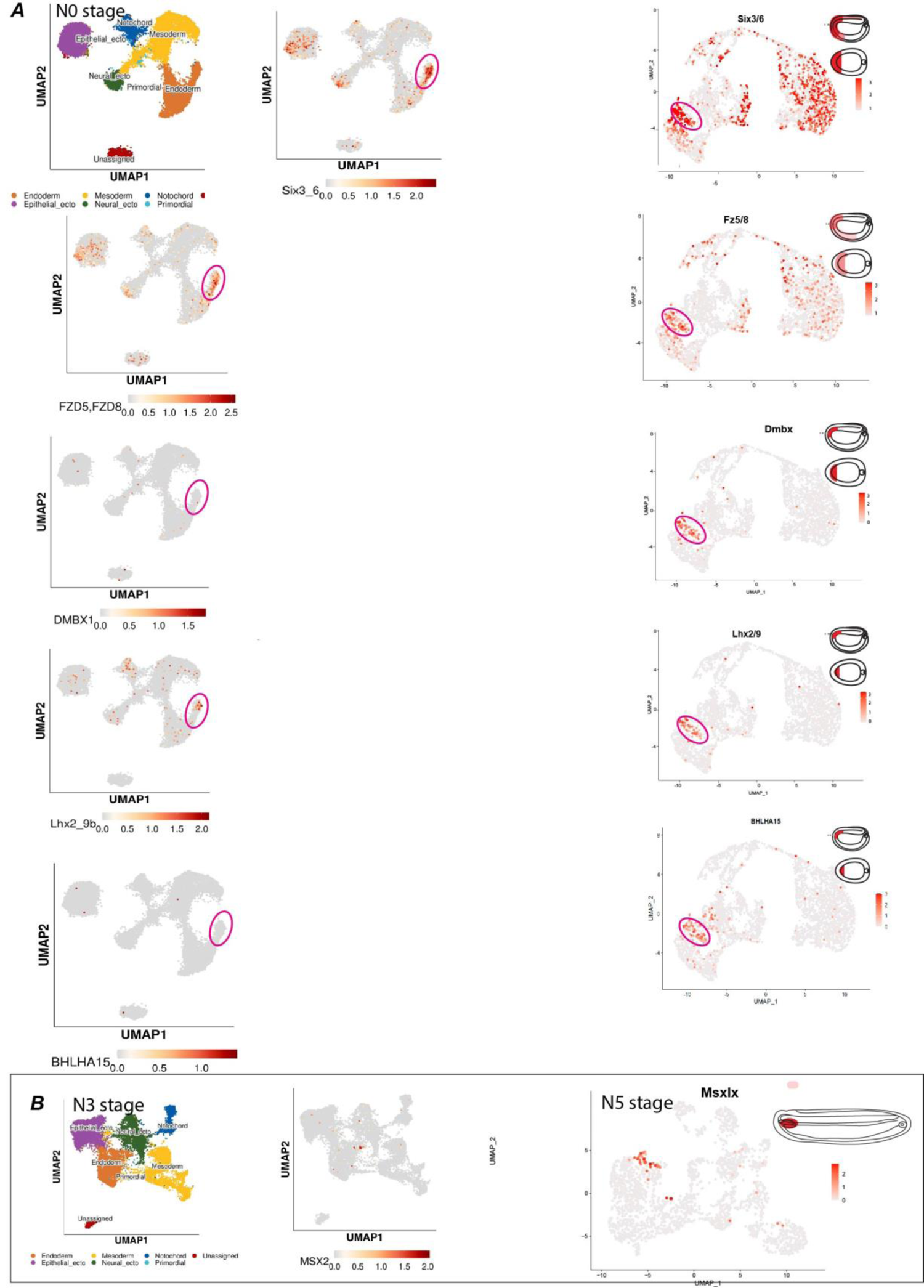
Comparison of presented scRNA-seq dataset with the dataset of Ma *et al.* Feature plots of PrCP-specific genes at N0 and Msxls at N3 and N5 in the dataset of Ma *et al.* (to the left) and the dataset presented here (to the right). Pink circle indicates presumed PrCP in the dataset of Ma *et al.* and PrCP in the dataset presented here.

**Table S1.**
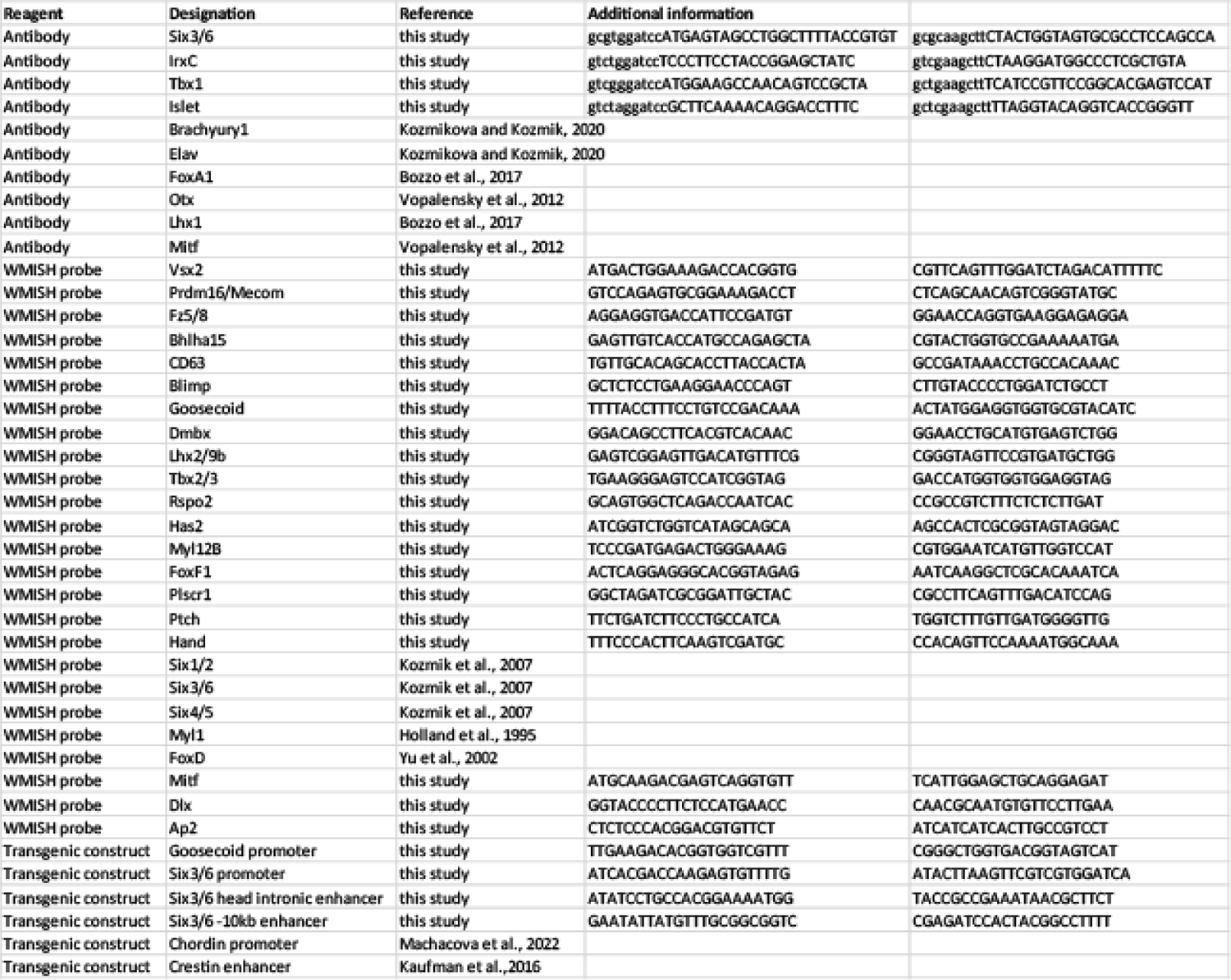

**Table S2.**

| sample | cells | median reads / cell | median genes / cell | reads |
| --- | --- | --- | --- | --- |
| G4 | 3,287 | 68,169 | 867 | 224,069,984 |
| N0 | 3,549 | 56,492 | 534 | 200,490,535 |
| N2 | 3,391 | 82,852 | 395 | 280,950,023 |
| N5 | 1,981 | 127,966 | 630 | 253,501,352 |

